# Differential expression of glutamate transporters and monoaminergic genes in major depression and suicide

**DOI:** 10.1101/596403

**Authors:** Brian Powers, Joel E. Kleinman, Thomas M. Hyde, Olusola Ajilore, Alex Leow, Monsheel S. Sodhi

**Affiliations:** Dept. Molecular Pharmacology and Therapeutics, Loyola University Chicago, Maywood, IL 60153; Lieber Institute for Brain Development and Department of Psychiatry & Behavioral Sciences, Johns Hopkins University School of Medicine, Baltimore, MD; Department of Neurology, Johns Hopkins University School of Medicine, Baltimore, MD; Dept. Psychiatry, University of Illinois at Chicago, Chicago, IL 60613

## Abstract

Accumulating evidence indicates that the glutamate and monoamine systems contribute to the pathophysiology of major depressive disorder (MDD) and suicide. We have tested the expression of genes encoding glutamate transporters and monoaminergic proteins in the dorsolateral prefrontal cortex (DLPFC) of MDD subjects who died by suicide (MDD-S, n=51), MDD non-suicide subjects (MDD-NS, n=28), and non-psychiatric controls (CTRL, n=32). We analyzed glutamate transporters (EAAT1, EAAT2, VGLUT1, and VGLUT2) and monoaminergic genes (SERT, NET, DAT, PMAT, VMAT, TPH1 and TPH2). Females but not males with MDD showed higher expression of all glutamate transporters relative to CTRLs (P<0.05). MDD-S groups of both sexes had higher VGLUT2 expression (P<0.05). MDD-S females who were antidepressant positive (+) had lower EAAT1 expression (P=0.004), perhaps indicating poor treatment response. Analyses of monoaminergic genes revealed lower VMAT1 expression (P=0.002) in MDD males, and conversely higher VMAT2 in MDD females (P=0.004). MDD females also had higher VMAT2, TPH2 and NET expression (p<0.05), and in contrast, MDD males had lower VMAT1 and PMAT expression. Therefore, we report sex differences in the expression of glutamate transporters and some monoaminergic genes in the DLPFC in MDD. Most of these findings are novel, but lower EAAT1 expression in MDD-S replicates previous studies. Lower EAAT1 expression coupled with higher VGLUT2 expression in MDD-S may lead to increased synaptic glutamate, neuronal loss and glial loss in the DLPFC in MDD and suicide reported previously. These deficits may contribute to lower DLPFC activity, poor problem solving and impaired executive function exhibited in severe depression and suicide.

## Introduction

Major depressive disorder (MDD) is a chronic psychiatric disorder that leads to significant medical and economic burdens on individuals and societies. It affects mood and behavior, resulting in cognitive impairments, sleep disturbances, alteration of appetite, anxiety, and anhedonia. MDD is also characterized by thoughts of suicide. Over 50% of individuals who die by suicide have a primary diagnosis of MDD (1, 2).

Conventional antidepressant treatments target the monoamine system(3–6), but only one-third of patients demonstrate substantial improvement, and many fail to respond to treatment (7–9). The commonly prescribed antidepressant drugs include selective serotonin reuptake inhibitors (SSRIs), which are antagonists of the serotonin (5-Hydroxytryptamine, 5-HT) transporter protein (SERT); serotonin-norepinephrine reuptake inhibitors (SNRIs), which inhibit both SERT and the norepinephrine transporter protein (NET); tricyclic antidepressants (TCAs), which increase synaptic release of monoamines; and monoamine oxidase inhibitors (MAOIs), which inhibit the metabolism of monoamines (10). Although these treatments rapidly increase the synaptic levels of 5-HT, norepinephrine and dopamine, there is usually a delay of several weeks before patients achieve antidepressant efficacy. In contrast, glutamatergic drugs, such as ketamine, have rapid antidepressant efficacy (11) in treatment resistant, severely depressed patients, sometimes within hours of treatment (12, 13). These and other accumulating data indicate glutamatergic abnormalities (14–16) and differences between the sexes in depression.

MDD is sexually dimorphic, with two to three fold greater frequency (17, 18), and greater symptom severity in women (19, 20). These clinical observations may reflect sex differences in molecular mechanisms underlying MDD(21). Transcriptional abnormalities in cortico-limbic brain regions that are associated with MDD differ greatly between the sexes. When genes are abnormally expressed in MDD in both sexes, these are often altered in opposite directions (22).

Brain regions implicated in the pathophysiology of MDD and suicide are sexually dimorphic (23). Among these regions, the dorsolateral prefrontal cortex (DLPFC) is critical for optimal emotional and executive function (24). During emotional tasks, the DLPFC is deactivated (25–27). Brain imaging studies show that executive function is disrupted in MDD patients (28) due to impaired DLPFC activity (29), perhaps due to reduced numbers of DLPFC synapses in MDD (30). Modulating DLPFC activity (31–36) may be necessary for antidepressant efficacy. The rapid antidepressant effects of ketamine, an antagonist of the N-methyl-D-aspartate receptor (NMDAR), coincide with rapid increases in spine synapses in the prefrontal cortex and activation of downstream signaling pathways that stimulate synapse formation (37). Accumulating data, including from our laboratory, indicate altered glutamatergic gene expression in the DLPFC in MDD (38–40).

There is a well-documented association between MDD and disrupted transmission within the glutamate(14–16) and monoamine systems (41–46). Although many of the conventional antidepressants target monoaminergic transmission, there are few studies reporting mRNA expression levels of monoaminergic genes in the DLPFC in MDD. In the current study, we have tested the hypothesis that glutamate transporter and monoaminergic gene expression are abnormal in the DLPFC of MDD patients, and that these gene expression abnormalities differ between the sexes. We have tested this hypothesis in a large cohort of postmortem subjects from three diagnostic groups: MDD patients who died by suicide (MDD-S), MDD patients who did not die by suicide (MDD-NS), and a comparison group of control subjects with no history of psychiatric illness (CTRL). Given that antidepressant treatment response may be predicted by suicidality in patients (47), we also tested if gene expression differs between the MDD-S and MDD-NS groups when assessing only subjects who are likely to have been compliant with antidepressant therapy at their time of death, as indicated by a positive test for antidepressants postmortem (MDD-S+ and MDD-NS+). Lastly, we have tested if any altered monoaminergic gene expression in the DLPFC results from antidepressant treatment, by comparing MDD patients who either did or did not test positive for antidepressants postmortem (MDD+ and MDD-). The results of this study indicate that glutamate transporter expression was upregulated in female but not male subjects with MDDrelative to CTRL groups. We also report sex-specific effects in the expression of some monoaminergic genes.

## Materials and Methods

### Subjects

Human postmortem brain specimens were obtained from the Brain Collection of the Clinical Brain Disorders Branch (subsequently renamed Human Brain Collection Core) at the National Institute of Mental Health (NIMH). Brains were collected from the Offices of the Chief Medical Examiner in Washington, DC and in Northern Virginia, USA. Postmortem interval (PMI) was calculated as time elapsed between death and tissue freezing, in hours. Frozen tissue from the DLPFC was obtained from two groups of subjects: (1) patients diagnosed with MDD, based on Diagnostic and Statistical Manual of Mental Disorders, Fourth Edition (DSM-IV) criteria; and (2) a control group with no history of substance abuse, psychiatric or neurological disorders. Each case had a 20-item telephone screening performed by a physician on the day of donation to document medical, psychiatric, substance abuse and social history. Psychiatric records and/or family informant interviews (computer-assisted Structured Clinical Interview for DSM-IV Disorders and NIMH Psychological Autopsy interview) were conducted with all psychiatric cases, when possible, by a master’s level clinician. Psychiatric narrative summaries were prepared, incorporating all of the above information. Two board-certified psychiatrists reviewed these cases to arrive at lifetime DSM-IV diagnoses. Table 1 summarizes the demographic variables of the postmortem subjects.

**Table 1.**
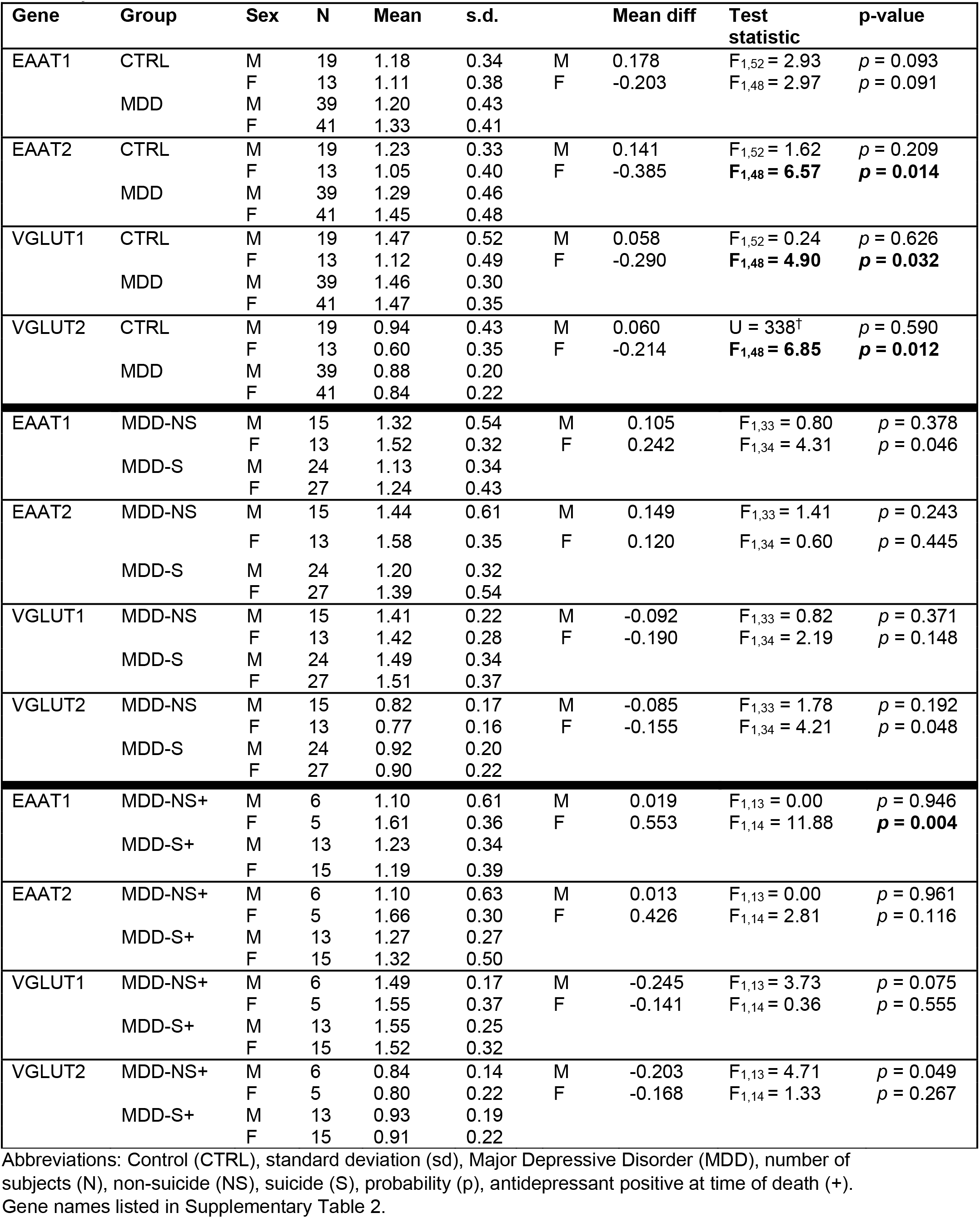
Expression of glutamate transporter genes in MDD and suicide. All analyses conducted in the DLPFC.

| Gene | Group | Sex | N | Mean | s.d. |  | Mean diff | Test statistic | p-value |
| --- | --- | --- | --- | --- | --- | --- | --- | --- | --- |
| EAAT1 | CTRL | M | 19 | 1.18 | 0.34 | M | 0.178 | $F_{1,52} = 2.93$ | $p = 0.093$ |
| | | F | 13 | 1.11 | 0.38 | F | -0.203 | $F_{1,48} = 2.97$ | $p = 0.091$ |
|  | MDD | M | 39 | 1.20 | 0.43 |  |  |  |  |
|  |  | F | 41 | 1.33 | 0.41 |  |  |  |  |
| EAAT2 | CTRL | M | 19 | 1.23 | 0.33 | M | 0.141 | $F_{1,52} = 1.62$ | $p = 0.209$ |
|  |  | F | 13 | 1.05 | 0.40 | F | -0.385 | **$F_{1,48} = 6.57$** | **$p = 0.014$** |
|  | MDD | M | 39 | 1.29 | 0.46 |  |  |  |  |
|  |  | F | 41 | 1.45 | 0.48 |  |  |  |  |
| VGLUT1 | CTRL | M | 19 | 1.47 | 0.52 | M | 0.058 | $F_{1,52} = 0.24$ | $p = 0.626$ |
|  |  | F | 13 | 1.12 | 0.49 | F | -0.290 | **$F_{1,48} = 4.90$** | **$p = 0.032$** |
|  | MDD | M | 39 | 1.46 | 0.30 |  |  |  |  |
|  |  | F | 41 | 1.47 | 0.35 |  |  |  |  |
| VGLUT2 | CTRL | M | 19 | 0.94 | 0.43 | M | 0.060 | $U = 338^{\dagger}$ | $p = 0.590$ |
|  |  | F | 13 | 0.60 | 0.35 | F | -0.214 | **$F_{1,48} = 6.85$** | **$p = 0.012$** |
|  | MDD | M | 39 | 0.88 | 0.20 |  |  |  |  |
|  |  | F | 41 | 0.84 | 0.22 |  |  |  |  |
| EAAT1 | MDD-NS | M | 15 | 1.32 | 0.54 | M | 0.105 | $F_{1,33} = 0.80$ | $p = 0.378$ |
| | | F | 13 | 1.52 | 0.32 | F | 0.242 | $F_{1,34} = 4.31$ | $p = 0.046$ |
|  | MDD-S | M | 24 | 1.13 | 0.34 |  |  |  |  |
|  |  | F | 27 | 1.24 | 0.43 |  |  |  |  |
| EAAT2 | MDD-NS | M | 15 | 1.44 | 0.61 | M | 0.149 | $F_{1,33} = 1.41$ | $p = 0.243$ |
| | | F | 13 | 1.58 | 0.35 | F | 0.120 | $F_{1,34} = 0.60$ | $p = 0.445$ |
|  | MDD-S | M | 24 | 1.20 | 0.32 |  |  |  |  |
|  |  | F | 27 | 1.39 | 0.54 |  |  |  |  |
| VGLUT1 | MDD-NS | M | 15 | 1.41 | 0.22 | M | -0.092 | $F_{1,33} = 0.82$ | $p = 0.371$ |
| | | F | 13 | 1.42 | 0.28 | F | -0.190 | $F_{1,34} = 2.19$ | $p = 0.148$ |
|  | MDD-S | M | 24 | 1.49 | 0.34 |  |  |  |  |
|  |  | F | 27 | 1.51 | 0.37 |  |  |  |  |
| VGLUT2 | MDD-NS | M | 15 | 0.82 | 0.17 | M | -0.085 | $F_{1,33} = 1.78$ | $p = 0.192$ |
| | | F | 13 | 0.77 | 0.16 | F | -0.155 | $F_{1,34} = 4.21$ | $p = 0.048$ |
|  | MDD-S | M | 24 | 0.92 | 0.20 |  |  |  |  |
|  |  | F | 27 | 0.90 | 0.22 |  |  |  |  |
| EAAT1 | MDD-NS+ | M | 6 | 1.10 | 0.61 | M | 0.019 | $F_{1,13} = 0.00$ | $p = 0.946$ |
| | | F | 5 | 1.61 | 0.36 | F | 0.553 | $F_{1,14} = 11.88$ | **$p = 0.004$** |
|  | MDD-S+ | M | 13 | 1.23 | 0.34 |  |  |  |  |
|  |  | F | 15 | 1.19 | 0.39 |  |  |  |  |
| EAAT2 | MDD-NS+ | M | 6 | 1.10 | 0.63 | M | 0.013 | $F_{1,13} = 0.00$ | $p = 0.961$ |
| | | F | 5 | 1.66 | 0.30 | F | 0.426 | $F_{1,14} = 2.81$ | $p = 0.116$ |
|  | MDD-S+ | M | 13 | 1.27 | 0.27 |  |  |  |  |
|  |  | F | 15 | 1.32 | 0.50 |  |  |  |  |
| VGLUT1 | MDD-NS+ | M | 6 | 1.49 | 0.17 | M | -0.245 | $F_{1,13} = 3.73$ | $p = 0.075$ |
| | | F | 5 | 1.55 | 0.37 | F | -0.141 | $F_{1,14} = 0.36$ | $p = 0.555$ |
|  | MDD-S+ | M | 13 | 1.55 | 0.25 |  |  |  |  |
|  |  | F | 15 | 1.52 | 0.32 |  |  |  |  |
| VGLUT2 | MDD-NS+ | M | 6 | 0.84 | 0.14 | M | -0.203 | $F_{1,13} = 4.71$ | $p = 0.049$ |
| | | F | 5 | 0.80 | 0.22 | F | -0.168 | $F_{1,14} = 1.33$ | $p = 0.267$ |
|  | MDD-S+ | M | 13 | 0.93 | 0.19 |  |  |  |  |
|  |  | F | 15 | 0.91 | 0.22 |  |  |  |  |

Abbreviations: Control (CTRL), standard deviation (sd), Major Depressive Disorder (MDD), number of subjects (N), non-suicide (NS), suicide (S), probability (p), antidepressant positive at time of death (+). Gene names listed in Supplementary Table 2.

The medical examiner’s office performed toxicological screenings in the blood, brain or other available matrix for prescribed medications, drugs of abuse and other substances that may have been related to the cause of death. Supplemental toxicology screening was done by the NIMH in order to fill in any gaps in testing for both prescription medications and drugs of abuse. Selective supplemental toxicology was performed depending upon the history of each patient. To measure the pH of the brain, 100 mg of homogenized frozen brain tissue was mixed with 1ml of cooled deionized water and pH was measured using a FiveEasy Plus pH meter (Mettler-Toledo LLC, Columbus, OH, USA). Autopsy information was reviewed, including cause of death, to exclude cases with hepatic or renal disease (which may cause increases in astroglia). Macroscopic and microscopic examination of the brain, including Bielschowsky’s silver stain (adapted for paraffin sections) on multiple cortical areas, was used to exclude cases with neuritic pathology, such as Alzheimer’s disease or cerebrovascular accidents. All layers of the cortex were removed. The middle frontal gyrus incorporating BA9/46 was dissected from a 1 cm thick slab just rostral to the rostrum of the corpus callosum, following the maps generated by Rajkowska and Goldman-Rakic (48).

### Preparation of cDNA

Laboratory personnel were blind to clinical data during the experiments. RNA was extracted from gray matter and complementary DNA (cDNA) prepared by standard methods (39). RNA integrity number (RIN), was measured using an Agilent BioAnalyzer (Agilent Technologies, Santa Clara, CA, USA). The average RIN value was 8.3, which indicates high quality RNA (Supplementary Table 1). Equal quantities of RNA from each subject were used to synthesize cDNA using the High Capacity cDNA Reverse Transcription Kit (Applied Biosystems, Foster City, CA, USA). Aliquots of cDNA from each subject were pooled for use as standards to generate a calibration curve according to the relative standard curve method (http://www.AppliedBiosystems.com). All RNA samples were diluted to 20 ng per microliter, followed by preamplification using the High Capacity cDNA Reverse Transcription Kit (Applied Biosystems). Preamplification of cDNA was necessary due to the low starting concentration of mRNA. Equal volumes of each TaqMan assay to be used for expression analysis were combined for the preamplification reaction. Forty microliter cDNA, together with 4 μl pooled assays and 44 μl TaqMan JumpStart (Sigma-Aldrich, St Louis, MO, USA), was preamplified for 14 cycles as previously described (49, 50).

### Gene expression assays

We measured gene expression as previously described (39). Assays are summarized in Supplementary Table 2. Quantitative PCR (QPCR) assays were performed in duplicate using the Applied Biosystems ViiA 7 System (Life Technologies, Grand Island, NY, USA). Housekeeping genes used to normalize the gene expression data were eliminated if associated with independent variables and covariates in analyses of variance (ANOVA). Of six housekeeping (HK) genes tested, three (GUSB, B2M, ACTB) were included in our analyses. The geometric mean of HK gene expression was not associated with diagnosis or sex, nor was there an interaction of sex by diagnosis. Normalizing data with multiple HK genes has greater accuracy than calculations from a single housekeeping gene. The relative expression levels of test transcripts were calculated using the Relative Standard Curve Method (www.AppliedBiosystems.com).

### Statistical analyses

We tested the hypothesis that dysfunction of glutamate and monoamine systems occurs in MDD and suicide by examining the expression of glutamate transporter genes and monoaminergic genes in the DLPFC of a postmortem cohort including patients with MDD (n=80) and a control group of psychiatrically healthy subjects (n=32). We used multivariate analysis of covariance (MANCOVA) to investigate main effects, and performed *posthoc* analyses of individual genes using univariate ANCOVA, with PMI, age at death, brain pH, and RIN included as covariates. We used Box’s M test to confirm that the observed covariance matrices for the dependent variables was equal across groups. We used Levene’s test of homogeneity of variance to confirm that the variance in each group compared was similar. If data that violated Levene’s test, we used the non-parametric Mann-Whitney U test for data analysis. We also tested the relationship between gene expression and demographic variables using analysis of variance (Supplementary Table 1). We calculated the false discovery rate (FDR) to correct for the effects of multiple comparisons (51). Statistical analyses were performed using IBM SPSS Statistics for Windows, Version 24.0. (IBM Corp., Armonk, NY, USA).

In light of recent reports, including our own, indicating dramatic sexual dimorphism in gene expression profiles in MDD (21, 22, 39), we analyzed males and females separately.

**For glutamate transporter gene expression analyses**, the following model was tested using MANCOVA:

1. Gene expression (y) = β0 + β1PMI + β2Age + β3pH + β4RIN + β5Diagnosis Diagnosis was a two-level factor, Control vs MDD.

Subsequently, we compared subgroups of the MDD cases to test for potential predictors of suicide, using the following model:

2. Gene expression (y) = β0 + β1PMI + β2Age + β3pH + β4RIN + β5Suicide Suicide was a two-level factor, MDD-NS vs MDD-S.

We then compared subgroups of the MDD patients who tested positive for antidepressants postmortem, and either did or did not die by suicide, using the model below:

3. Gene expression (y) = β0 + β1PMI + β2Age + β3pH + β4RIN + β5Suicide Suicide was a two-level factor, MDD-NS+ vs MDD-S+.

**For monoamine gene expression analysis**, the following model was tested using MANCOVA:

4. Gene expression (y) = β0 + β1PMI + β2Age + β3pH + β4RIN + β5Diagnosis Diagnosis was a two-level factor, Control vs MDD.

We then compared subgroups of the MDD cases to test for potential predictors of suicide, using the following model:

5. Gene expression (y) = β0 + β1PMI + β2Age + β3pH + β4RIN + β5Suicide Suicide was a two-level factor, MDD-S vs MDD-NS.

We then compared subgroups of the MDD patients who tested positive for antidepressants postmortem, and either did or did not die by suicide, using the model below:

6. Gene expression (y) = β0 + β1PMI + β2Age + β3pH + β4RIN + β5Suicide Suicide was a two-level factor, MDD-S+ vs MDD-NS+.

Lastly, we compared monoaminergic gene expression between MDD patients who either did or did not test positive for antidepressants postmortem, using the following model:

7. Gene expression (y) = β0 + β1PMI + β2Age + β3pH + β4RIN + β5Antidepressant Antidepressant was a two-level factor, MDD+ vs MDD-.

## Results

Our multivariate analyses revealed increased expression of glutamate transporters in female patients diagnosed with MDD relative to controls (F_4,45_ = 2.77, P= 0.038). *Posthoc* univariate analyses indicated that EAAT2, VGLUT1, and VGLUT2 genes exhibited significantly increased expression in MDD subjects relative to controls (Figure 1A, Table 1A). In contrast, we observed no significant differences in glutamatergic gene expression in the male groups.

**Figure 1:**
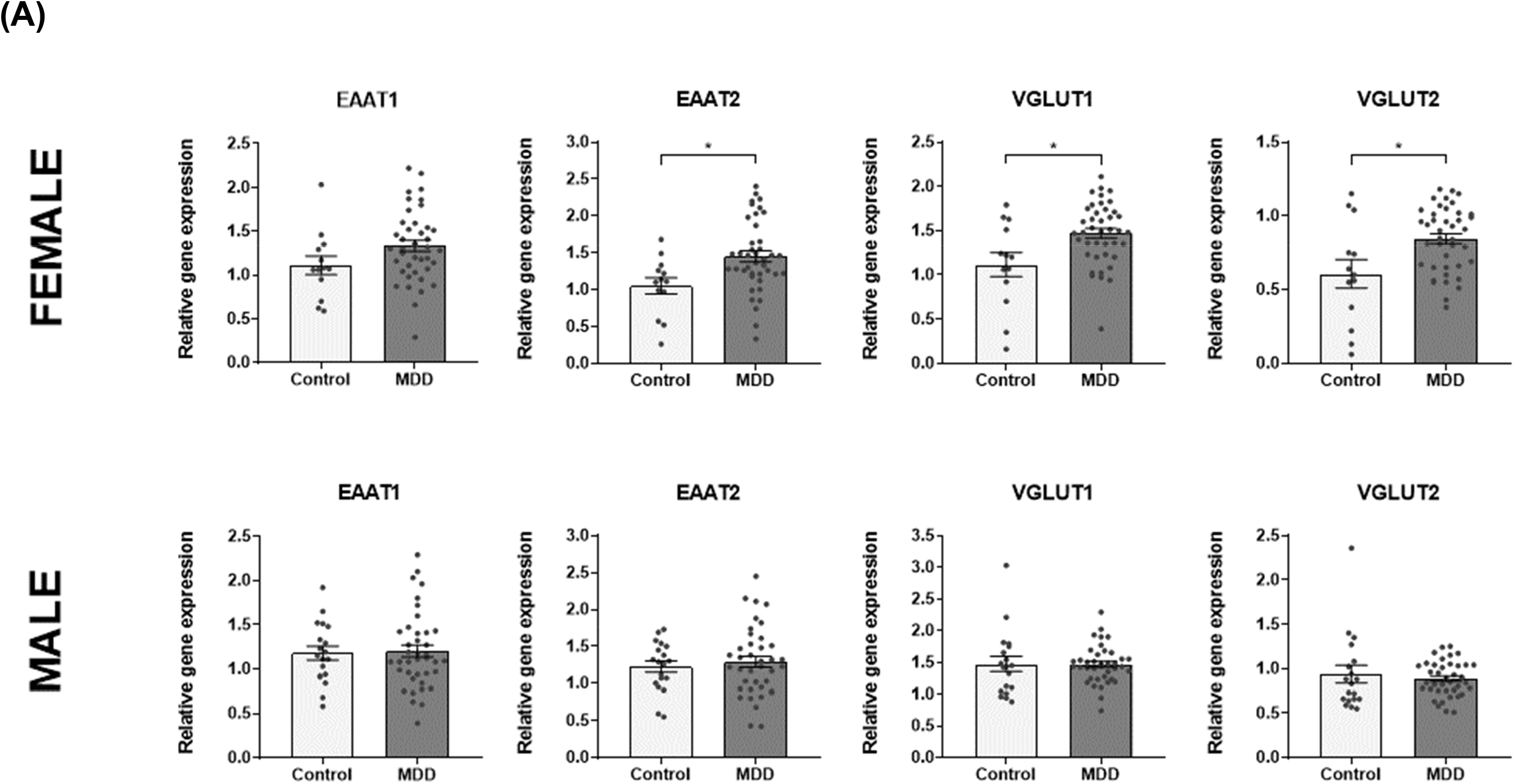
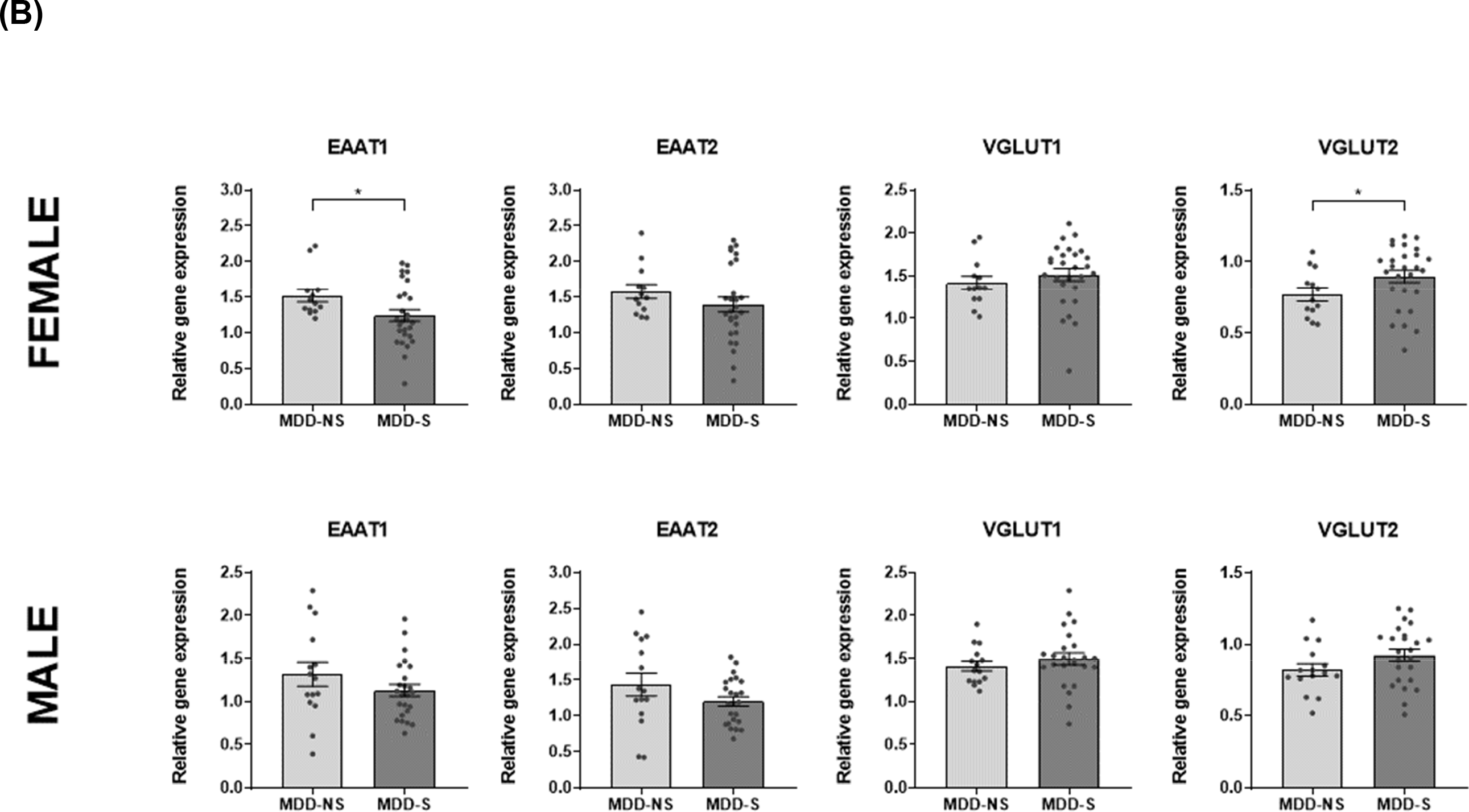
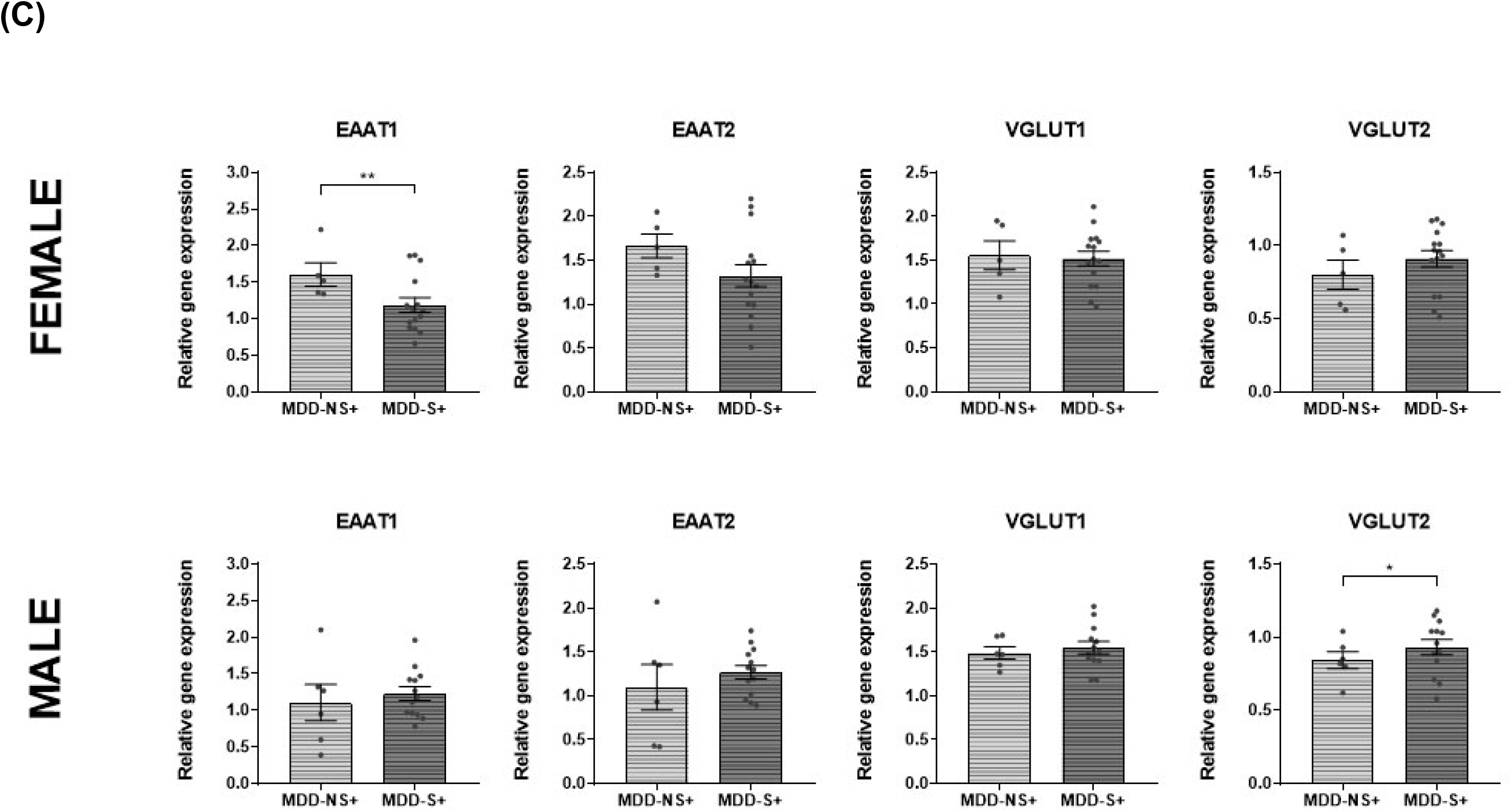
Sex differences in the expression of glutamate transporter genes in depression and suicide. (A) Elevated expression of glutamate transporter genes in depressed females. (B). Lower expression of EAAT1 and higher expression of VGLUT2 in depressed female suicides. (C). Suicide may be an indicator of antidepressant treatment response. Tests of depressed subjects who were antidepressant positive at time of death reveal that female suicides had lower expression of EAAT1 relative to the female non-suicides. In contrast, male suicides who were antidepressant positive, had slightly higher levels of VGLUT2 than male non-suicides. *p<0.05, ** p<0.01.

We also sought to determine potential predictors of suicide risk by analyzing gene expression in MDD suicides (MDD-S) compared to MDD non-suicides (MDD-NS). In female subjects, multivariate analysis revealed a significant difference in glutamate transporter gene expression between MDD-S and MDD-NS groups (F_4,31_ = 3.87, P= 0.012). *Posthoc* univariate analyses (Figure 1B, Table 1A) showed lower expression of EAAT1 and higher expression of VGLUT2 in the female MDD-S group compared to the female MDD-NS group (F_1,76_ =4.60, P= 0.035). There were no significant differences in glutamate transporter expression between the male MDD-S and MDD-NS subjects.

An additional analysis of potential predictors of treatment response included only MDD patients who tested positive for antidepressants postmortem. In female subjects, multivariate analysis showed a significant difference in glutamate transporter gene expression between MDD-S+ and MDD-NS+ groups (F_4,11_ = 5.07, P= 0.015). *Posthoc* univariate analyses revealed lower expression levels of EAAT1 in the female MDD-S+ group compared to the female MDD-NS+ group (Figure 1C, Table 1B). In male subjects, multivariate analyses of glutamate transporter expression showed no differences between the MDD-S+ and MDD-NS+ groups. However, *post-hoc* univariate analyses indicated increased expression of VGLUT2 in the MDD-S+ group compared to the MDD-NS+ group.

Subsequent analyses focused on the expression of monoaminergic genes. Multivariate analyses showed a significant effect of diagnosis in female subjects (F_8,41_ = 2.41, P= 0.031) and male subjects (F_8,45_ = 2.23, P= 0.043). *Post-hoc* univariate analyses revealed increased expression of VMAT2, NET, and TPH2 in females with MDD relative to controls (Figure 2A, Table 2). In contrast, univariate analyses indicate that males with MDD have decreased expression of VMAT1 and decreased expression of PMAT relative to controls (Figure 2B, Table 2).

**Figure 2:**
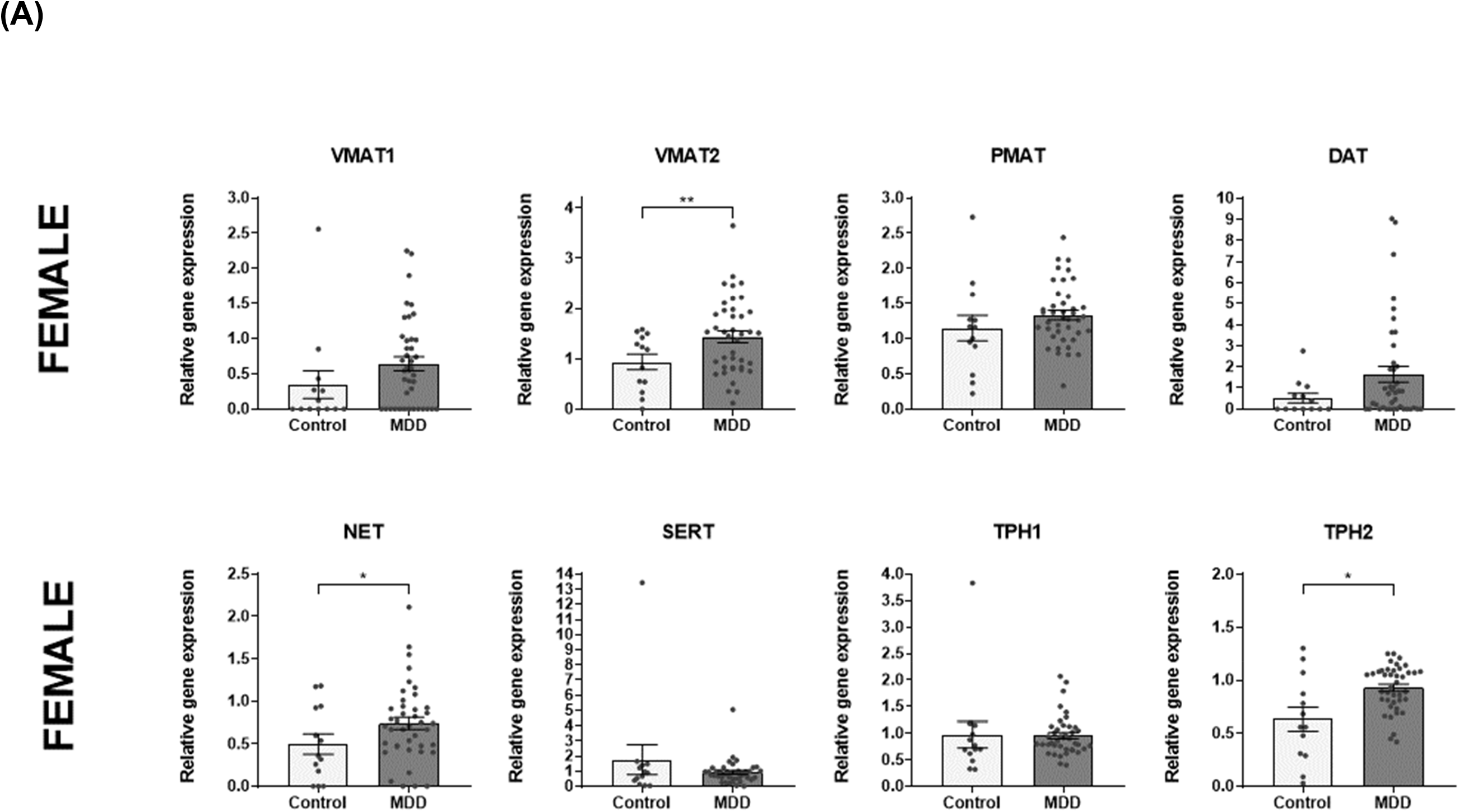
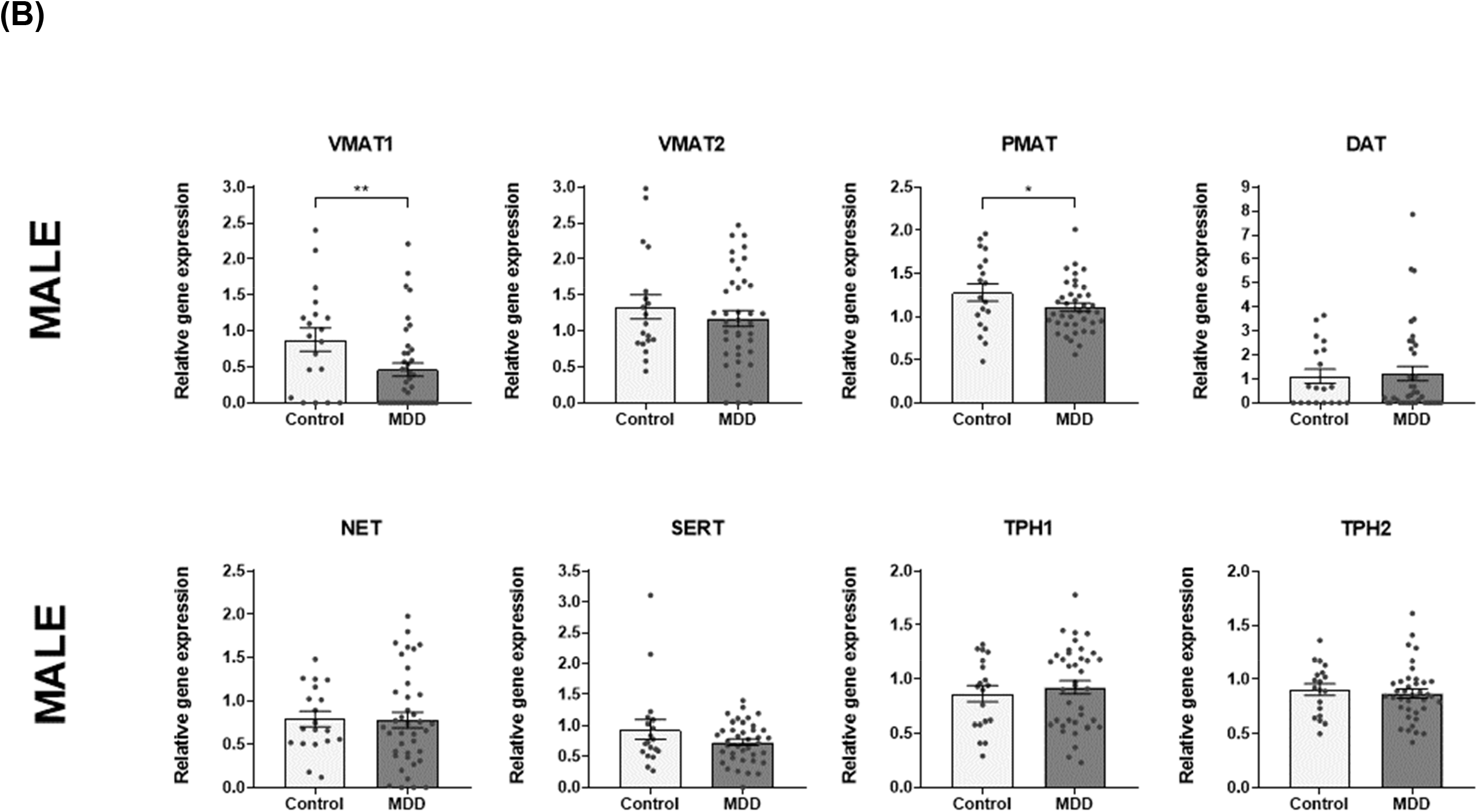
Sex differences in the expression of monoaminergic genes in depression and suicide. (A) Higher expression of VMAT2, NET and TPH2 in depressed females relative to female controls. (B). Lower expression of VMAT1 and PMAT in depressed males relative to male controls. *p<0.05, ** p<0.01.

**Table 2.** Sex differences in monoamine gene expression in depression. Depressed males had lower expression of VMAT1 relative to male controls, while depressed females had higher expression of VMAT2, NET and TPH2 relative to female controls. All analyses conducted in the DLPFC.

| Gene | Group | Sex | N | Mean | s.d. | Mean diff | statistic | p-value |
| --- | --- | --- | --- | --- | --- | --- | --- | --- |
| VMAT1 | CTRL | M | 19 | 0.88 | 0.71 | M 0.586 | $F_{1,52} = 10.62$ | **$p = 0.002$** |
| | | F | 13 | 0.34 | 0.71 | F -0.263 | $F_{1,48} = 1.59$ | $p = 0.214$ |
|  | MDD | M | 39 | 0.46 | 0.58 |  |  |  |
|  |  | F | 41 | 0.64 | 0.63 |  |  |  |
| VMAT2 | CTRL | M | 19 | 1.33 | 0.73 | M 0.211 | $F_{1,52} = 0.97$ | $p = 0.329$ |
| | | F | 13 | 0.94 | 0.56 | F -0.673 | $F_{1,48} = 8.92$ | **$p = 0.004$** |
|  | MDD | M | 39 | 1.17 | 0.67 |  |  |  |
|  |  | F | 41 | 1.43 | 0.75 |  |  |  |
| PMAT | CTRL | M | 19 | 1.28 | 0.44 | M 0.194 | $F_{1,52} = 3.97$ | $p = 0.050$ |
| | | F | 13 | 1.15 | 0.66 | F -0.250 | $F_{1,48} = 2.86$ | $p = 0.097$ |
|  | MDD | M | 39 | 1.11 | 0.30 |  |  |  |
|  |  | F | 41 | 1.33 | 0.44 |  |  |  |
| DAT | CTRL | M | 19 | 1.10 | 1.29 | M 0.445 | $F_{1,52} = 0.89$ | $p = 0.351$ |
| | | F | 13 | 0.51 | 0.80 | F -0.635 | $U = 184^{\dagger}$ | $p = 0.088$ |
|  | MDD | M | 39 | 1.22 | 1.85 |  |  |  |
|  |  | F | 41 | 1.65 | 2.40 |  |  |  |
| NET | CTRL | M | 19 | 0.79 | 0.38 | M -0.033 | $F_{1,52} = 0.04$ | $p = 0.834$ |
| | | F | 13 | 0.50 | 0.43 | F -0.325 | $F_{1,48} = 6.32$ | **$p = 0.015$** |
|  | MDD | M | 39 | 0.78 | 0.55 |  |  |  |
|  |  | F | 41 | 0.74 | 0.44 |  |  |  |
| SERT | CTRL | M | 19 | 0.93 | 0.67 | M 0.220 | $F_{1,52} = 2.55$ | $p = 0.117$ |
| | | F | 13 | 1.76 | 3.55 | F 0.659 | $F_{1,48} = 1.23$ | $p = 0.272$ |
|  | MDD | M | 39 | 0.73 | 0.33 |  |  |  |
|  |  | F | 41 | 0.92 | 0.79 |  |  |  |
| TPH1 | CTRL | M | 19 | 0.86 | 0.33 | M -0.046 | $F_{1,52} = 0.17$ | $p = 0.678$ |
| | | F | 13 | 0.97 | 0.91 | F -0.029 | $F_{1,48} = 0.03$ | $p = 0.867$ |
|  | MDD | M | 39 | 0.92 | 0.37 |  |  |  |
|  |  | F | 41 | 0.96 | 0.37 |  |  |  |
| TPH2 | CTRL | M | 19 | 0.90 | 0.23 | M 0.007 | $F_{1,52} = 0.01$ | $p = 0.919$ |
|  |  | F | 13 | 0.63 | 0.40 |  |  |  |
| | MDD | M | 39 | 0.87 | 0.25 | F -0.307 | $U = 144^{\dagger}$ | **$p = 0.013$** |
|  |  | F | 41 | 0.93 | 0.21 |  |  |  |

Abbreviations: Control (CTRL), standard deviation (sd), Major Depressive Disorder (MDD), number of subjects (N), probability (p). <sup>†</sup>Mann-Whitney U Test. Gene names listed in Supplementary Table 1.

Further analyses of monoaminergic gene expression revealed no significant differences between MDD-S and MDD-NS groups in either males or females (Supplementary Figure 1A, Supplementary Figure 1B, and Supplementary Table 3) regardless of antidepressant positive status (Supplementary Figure 1C, Supplementary Figure 1D, Supplementary Table 4). Moreover, analyses revealed no differences in monoaminergic gene expression when we compared MDD+ with MDD- patients in groups of either sex (Supplementary Figure 1E, Supplementary Figure 1F, and Supplementary Table 5).

Some of the data that we report here replicates previous studies, as outlined in the Discussion. When we correct these analyses for false discovery using the method proposed by Benjamini and Hochberg (51), these findings do not remain statistically significant. However, we note that 12 of the 88 *post-hoc* tests (14%) show an uncorrected p value ≤0.05, which is approximately three times the rate of positives expected with an α (false positive) threshold of 5%.

## Discussion

Our findings support the hypothesis that both glutamatergic and monoaminergic genes have abnormal expression in the DLPFC in MDD and suicide. We observed sex differences in these gene expression measures when comparing MDD with CTRLs, and when we compared MDD-S with MDD-NS groups.

### Sex differences in glutamate transporter expression in MDD

MDD females but not MDD males had higher expression levels of glutamate transporter genes compared with CTRLs of the same sex. These data add to the accumulating evidence that there are glutamatergic abnormalities in MDD and suicide, particularly in females (39). Upregulation of VGLUTs may lead to the elevated levels of synaptic or extracellular glutamate observed in the DLPFC in MDD (52). Elevated glutamate levels may result from increased activity at the glutamate synapse in MDD females. These findings are consistent with our previous report of elevated glutamate receptor expression in MDD females(39). In females, estrogen activates estrogen receptors (ER), which bind to estrogen response elements (EREs) upstream of the promoters of specific genes in the prefrontal cortex (53). EREs are located immediately upstream of the EAAT2 gene (54) therefore upregulated EAAT2 expression may result from elevated estrogen signaling during stress. The increased expression of EAAT proteins in MDD females may be a compensatory mechanism by which excess glutamate is removed from the synapse during stressful life events (55, 56).

Previous studies of mixed-sex cohorts, including mostly male subjects, show decreased expression of glutamate transporters in MDD in the frontal cortex (38, 57–60). In contrast, we detected upregulated expression of glutamate transporters in the DLPFC of MDD females but not in MDD males. However, increased expression of synapse-related genes in the DLPFC of MDD females has been reported previously (22).

### Glutamate transporter expression is abnormal in depressed suicides

Sex differences in gene expression in MDD may underpin the sex differences in symptoms and severity of MDD (21, 22). Completed suicide may be an indicator of more severe illness in MDD. Male and female MDD-S groups showed similar abnormalities of glutamate transporter expression relative to same sex MDD-NS groups, although we detected larger differences in the females. Increased expression of VGLUT2 in MDD-S subjects, may predict increased release of glutamate at the synapse. In addition, EAAT1 expression was relatively lower in MDD-S patients. Other studies of mixed-sex cohorts have reported decreased expression of EAAT1 and EAAT2 in the DLPFC in MDD-S(58), and in suicides with schizophrenia (61) with smaller effects in males. These data are also consistent with a recent study of mostly male subjects (40). Moreover, previous reports indicate lower levels of EAAT1 and EAAT2 expression in MDD in the frontal cortex (38, 57–59), the hippocampus (62) and locus coeruleus (63). In combination, these data indicate lower EAAT expression in the DLPFC is associated with severe depression, in both sexes.

Expression of EAAT1 is primarily in the astrocyte in the DLPFC (55, 56, 64) and therefore, lower EAAT1 expression may indicate a relative loss of glia or glial function in the DLPFC of the MDD-S females. Lower EAAT activity would reduce the efficiency of removal of extracellular glutamate. Figure 3 summarizes the functions of the glutamate transporters and their relative roles in regulating synaptic glutamate levels. In combination, if MDD-S groups have reduced EAAT1 expression and increased VGLUT2 expression, excess extracellular and synaptic glutamate could result, leading to excitotoxic cell death in the DLPFC (65, 66). If validated, these findings could explain the reduced DLPFC activity observed in suicidal patients. Reduced activity in the DLPFC would lead to poor problem solving and poor planning (67), which may result in suicide attempts in response to stress.

**Figure 3:**
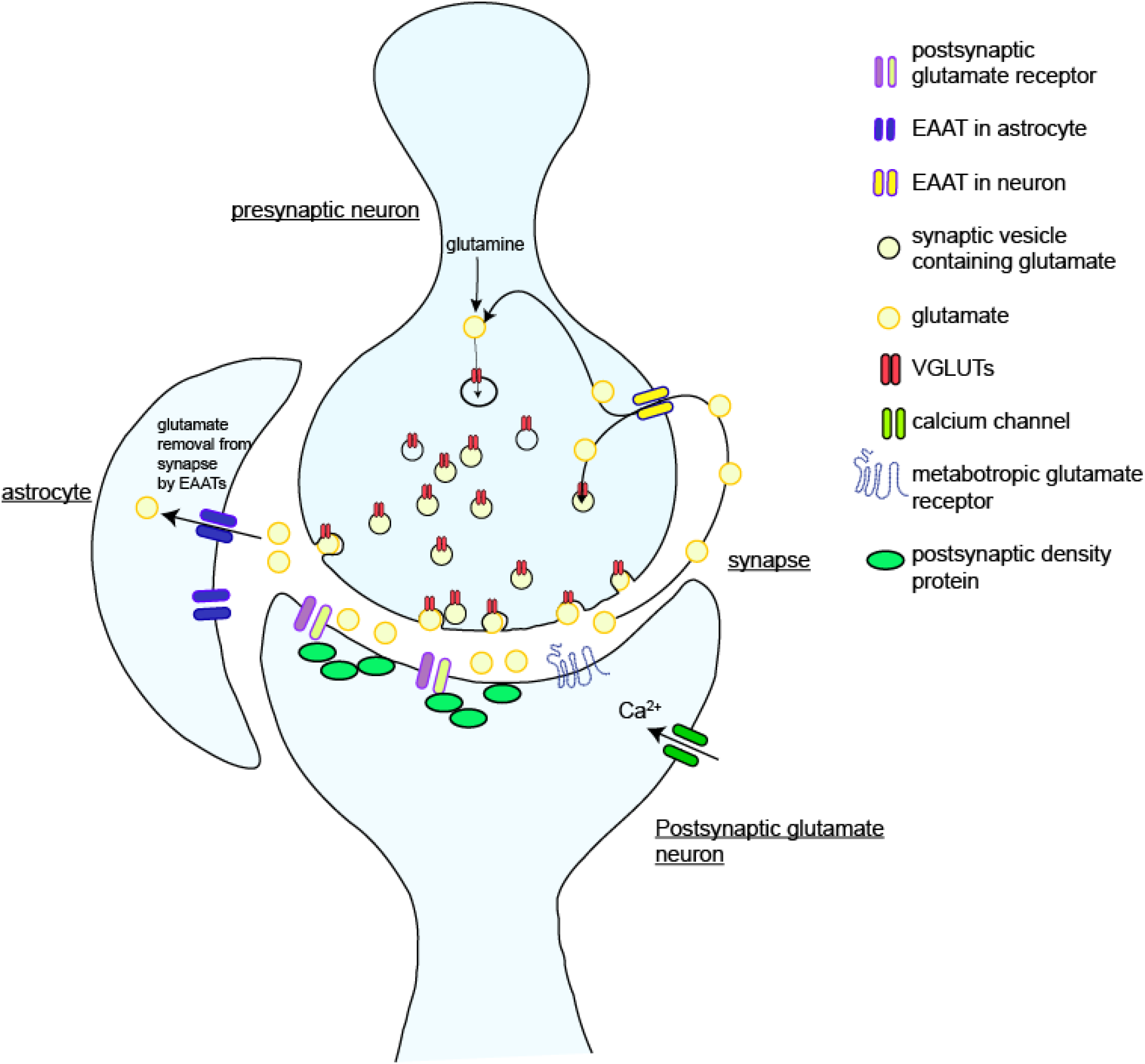
Illustration of glutamate transporters within the glutamate synapse. The three cells within the glutamatergic synapse include the presynaptic neuron, which expresses vesicular transporters (VGLUTs), the postsynaptic neuron, which expresses the majority of glutamate receptors, and the astrocytic end foot. The diagram illustrates the roles and locations of the glutamate transporters without details of postsynaptic density proteins and other molecular pathways. High affinity membrane transporters of glutamate in the glia (EAATs) are responsible for more than 90% of synaptic glutamate reuptake (81) and are essential for reducing synaptic glutamate concentrations. The high concentration of EAATs around synapses ensures that unbound glutamate will encounter available transporters (82). Reduced EAAT protein levels, lead to elevated extracellular glutamate, which leads to increased excitotoxicity and neuronal death (81). One model of MDD suggests decreased synaptic glutamate uptake and a consecutive elevation of extracellular glutamate levels in depression (83). This model and our findings are consistent with reductions in neuronal and glial density observed in post-mortem studies of MDD (84–87). Reduced glutamatergic activity in the DLPFC may explain hypoactivation of this brain region during cognitive and emotional processing tasks in MDD (88–94).

### Sex differences in glutamate transporter expression in antidepressant treatment responders

We have considered suicide completion in antidepressant positive MDD patients as an indication of poor response to antidepressant treatment. We compared the ‘poor responder group’ (MDD-S+) with a group of MDD patients who were antidepressant positive but who did not die by suicide i.e. ‘good responders’ (MDD-NS+). In females, we detected lower EAAT1 expression in MDD-S+ but in male MDD-S+, we detected higher VGLUT2 expression relative to MDD-NS+ of the same sex. Reduced EAAT1 or increased VGLUT2 would lead to higher glutamate levels in the synapse of both sexes (65). One previous report showed higher levels of synaptic glutamate in the DLPFC of MDD patients with poor treatment response (68). Therefore, males and females who respond poorly to antidepressants may have similarly elevated glutamate levels in the DLPFC but by slightly different mechanisms i.e. males may have greater glutamate release from the presynaptic neuron while females may have less efficient reuptake of extracellular glutamate. Therefore, our data indicate that different molecular or cellular abnormalities in males and females may underlie this abnormality of glutamate system regulation. One previous report indicates that female rats have enhanced sensitivity to ketamine during proestrus stage of the menstrual cycle, which is mediated by estradiol activation of the estrogen receptors, leading to greater activation of synaptic plasticity related kinases within prefrontal cortex (69).

### Sex differences in monoamine transporter expression in depression

Our expression analyses of monoaminergic genes included genes encoding the monoamine transporters and two isoenzymes of tryptophan hydroxylase. Considerable evidence implicates these genes in the pathophysiology of MDD and suicide (70) and several are targets of antidepressant drugs(71). We detected lower expression of VMAT1 and PMAT in MDD males compared with male controls. In the presynaptic terminals of monoaminergic neurons, vesicular monoamine transporters (VMATs) sequester 5-HT, dopamine, and norepinephrine into synaptic vesicles for release, which may be reduced if VMAT1 expression is lower in MDD males. PMAT is membrane bound and has a strong kinetic preference for 5-HT and dopamine although PMAT also transports other mono-amines, such as histamine, norepinephrine, and epinephrine(72). Lower PMAT expression in MDD males could indicate that there is reduced transport of several monoamines from the synapse. These altered expression levels are unlikely to be a result of antidepressant treatment received by the MDD patients, because we observed no differences in the expression of any monoaminergic gene between the antidepressant positive and antidepressant negative MDD patients.

In contrast with the MDD males, we observed increased expression of VMAT2, NET, and TPH2, in MDD females relative to controls. Comparison of antidepressant positive and antidepressant negative MDD female subjects indicates that the monoaminergic gene expression differences that we observe in the females are unlikely to be a result of antidepressant treatment. It is possible that some of these molecular abnormalities may predict antidepressant efficacy. Women show better response to SSRIs and MAOIs, and men have better response to TCAs (22). SSRIs increase synaptic 5-HT levels, while MAOIs are non-specific inhibitors of monoamine metabolism(10). The current data indicate disruption of gene expression in 5-HT neurons (due to TPH2 expression) and norepinephrine neurons (due to NET expression) in the DLPFC in women and perhaps drugs that target these systems, such as SSRIs and SNRIs will have greater antidepressant efficacy in females with MDD. In males, altered VMAT1 and PMAT expression indicates abnormalities that may not occur in any specific monoaminergic neuron(72) in MDD, which may be why non-specific monoaminergic drugs such as TCAs will have greater efficacy in males. While we observed no variation of DAT expression between our diagnostic groups, we cannot rule out abnormal dopaminergic transmission in MDD because we have observed altered expression of non-specific transporters such as the VMATs and PMAT in MDD. Moreover, VGLUT2, which was elevated in the suicide groups, is expressed in both glutamate and dopamine neurons (73). The upregulation of NET and TPH2 in MDD females may be due to stress-induced stimulation of estrogen response elements (EREs) immediately upstream of the NET, and TPH2 genes (54).

Our data may be consistent with a report of increased TPH2 mRNA expression in the prefrontal cortex in suicide (74), although there have been conflicting data (75). The expression of these genes was not associated with antidepressant positive status, indicating that these findings may not be influenced by the medication administered to these patients. We observed no abnormalities of SERT expression in MDD, which is consistent with a meta-analysis of five postmortem studies of the prefrontal cortex (76). To our knowledge, we are the first to report abnormal expression of the other VMAT, PMAT and NET in the DLPFC in MDD. We did not observe differences in monoaminergic gene expression in any other analysis.

*Limitations of this study* include those that are common to all postmortem analyses of human brain, but investigations of human postmortem brain are necessary for detailed cellular and molecular studies of neurological disorders. Demographic variables (Supplementary Table 1) including RIN values, sample number and detailed toxicological analyses performed are the strengths of this study (77). We based this study on strong neurobiological hypotheses, which is probably why we report three times the number of positive findings than would be expected by chance alone. Nevertheless, when we corrected our data for false discovery, the findings are no longer significant. Sole reliance on probability values to determine significance may be erroneous (78), and replication is a more reliable indicator of true discoveries. Replication is unusual when investigating human subjects, but in spite of this, some data that we report here replicate previous studies. The novel findings that we report will require replication to separate the true positives from the false positive data. In addition, mRNA abundance does not always correlate with protein activity. If some of the mRNA measured included non-coding RNAs, then altered mRNA levels would be less likely to correlate directly with protein activity.

Therefore, with the limitations we note in mind, we report sex differences in the expression of glutamate transporters and some monoaminergic genes in the DLPFC in MDD. Most of these findings are novel, but lower EAAT1 expression in MDD-S replicates previous studies. It should be noted that replication is rare in studies of human subjects. Lower EAAT1 expression coupled with higher VGLUT2 expression in MDD-S may lead to increased synaptic glutamate, neuronal loss and glial loss in the DLPFC in MDD and suicide reported previously. These deficits may contribute to lower DLPFC activity, poor problem solving and impaired executive function exhibited in severe depression and suicide. Taken together, these data highlight the importance of considering sex when investigating the biological basis of MDD and suicide. Future analyses of additional regions within the cortico-limbic circuitry in MDD would further elucidate the pathophysiological mechanisms of depression and suicide (79, 80).

## Acknowledgements and disclosures

Funded by the Hans W. Vahlteich Scholar Award (MS) and American Foundation for Suicide Prevention (MS). We thank the Human Brain Collection Core at NIMH. We also thank William Adams Ph.D. at the Biostatistics Core of Loyola University, for statistical advice. The authors report no conflicts of interest.

---

**Supplementary Figure 1:**
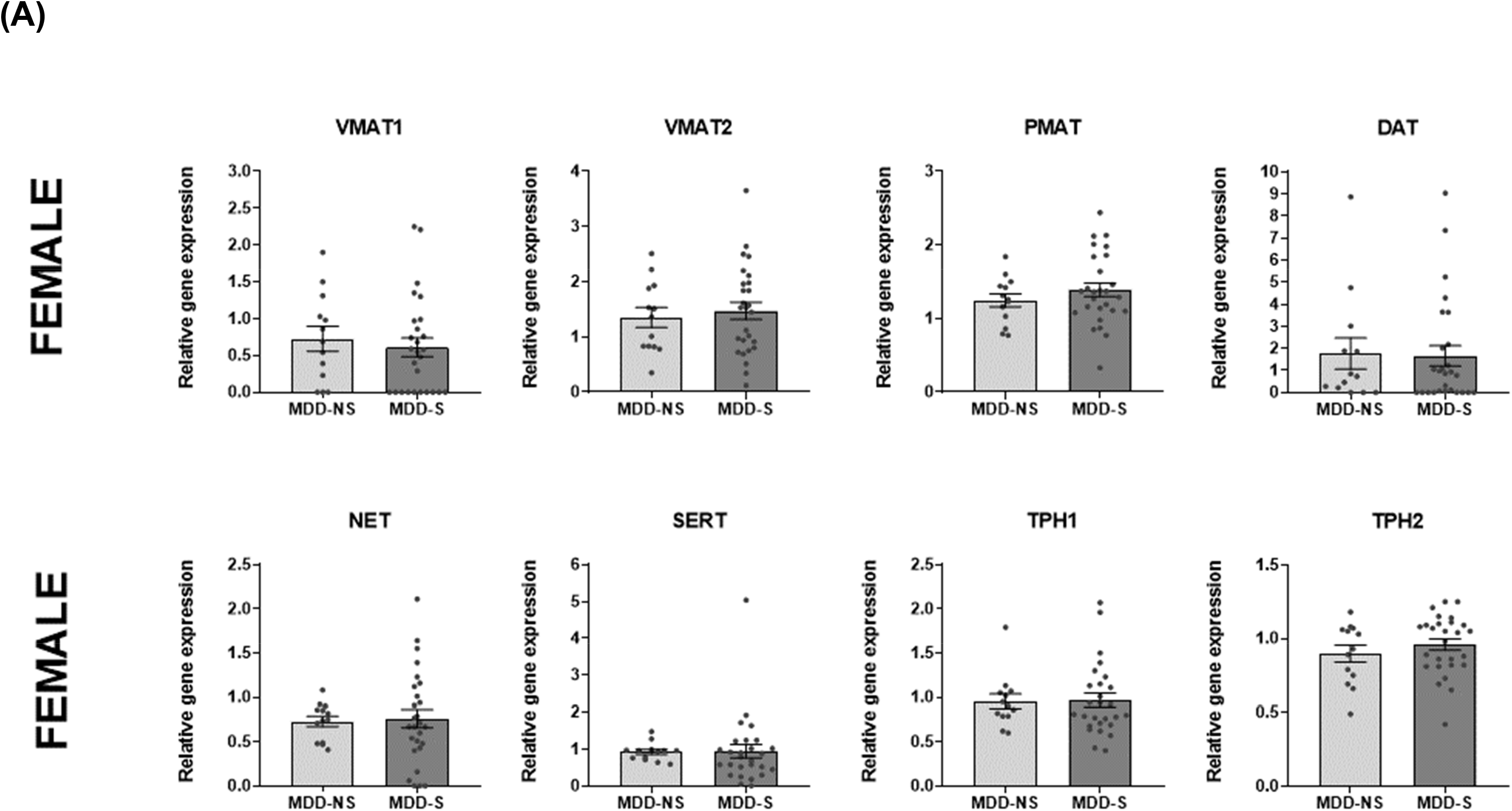
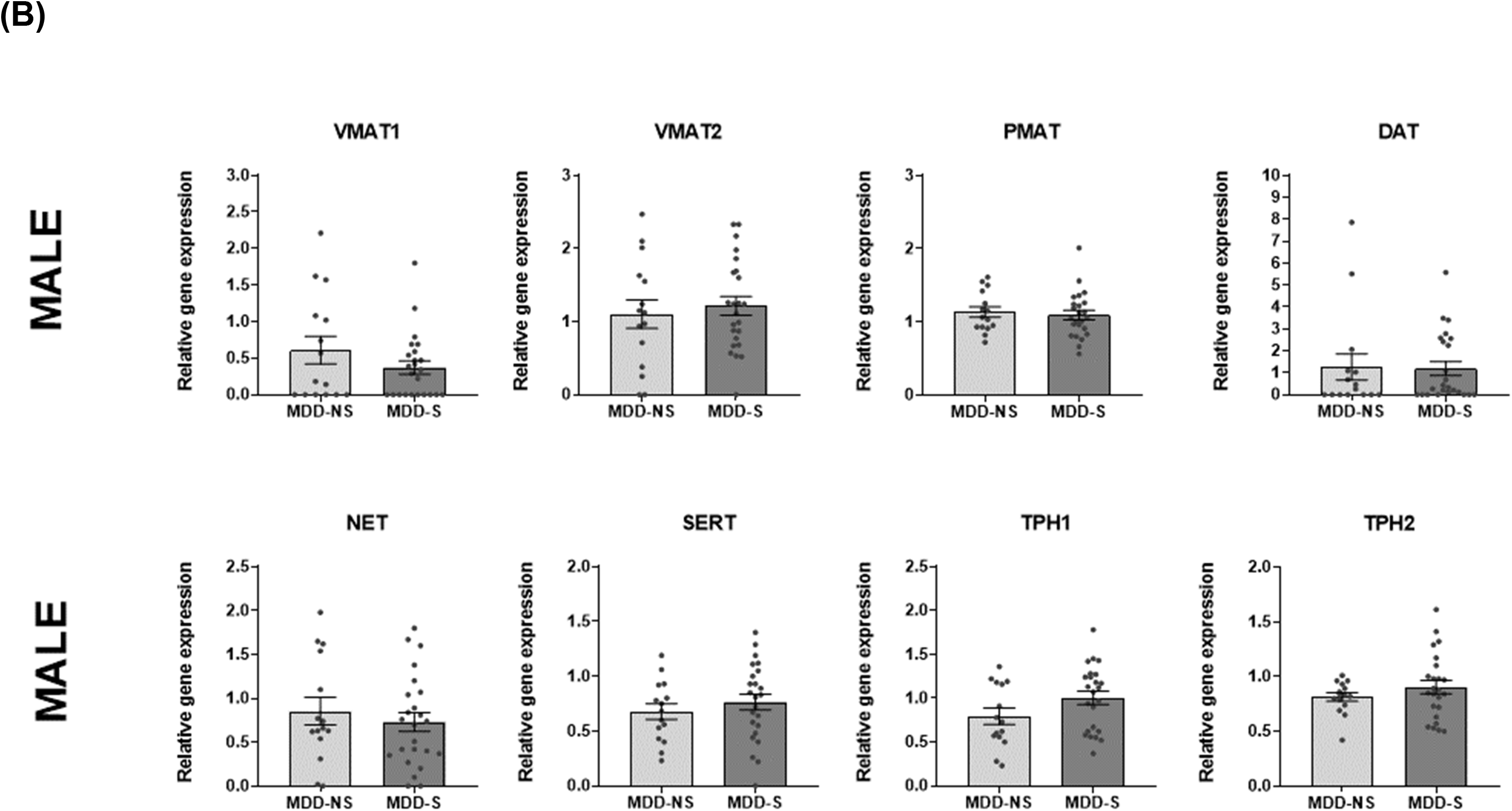
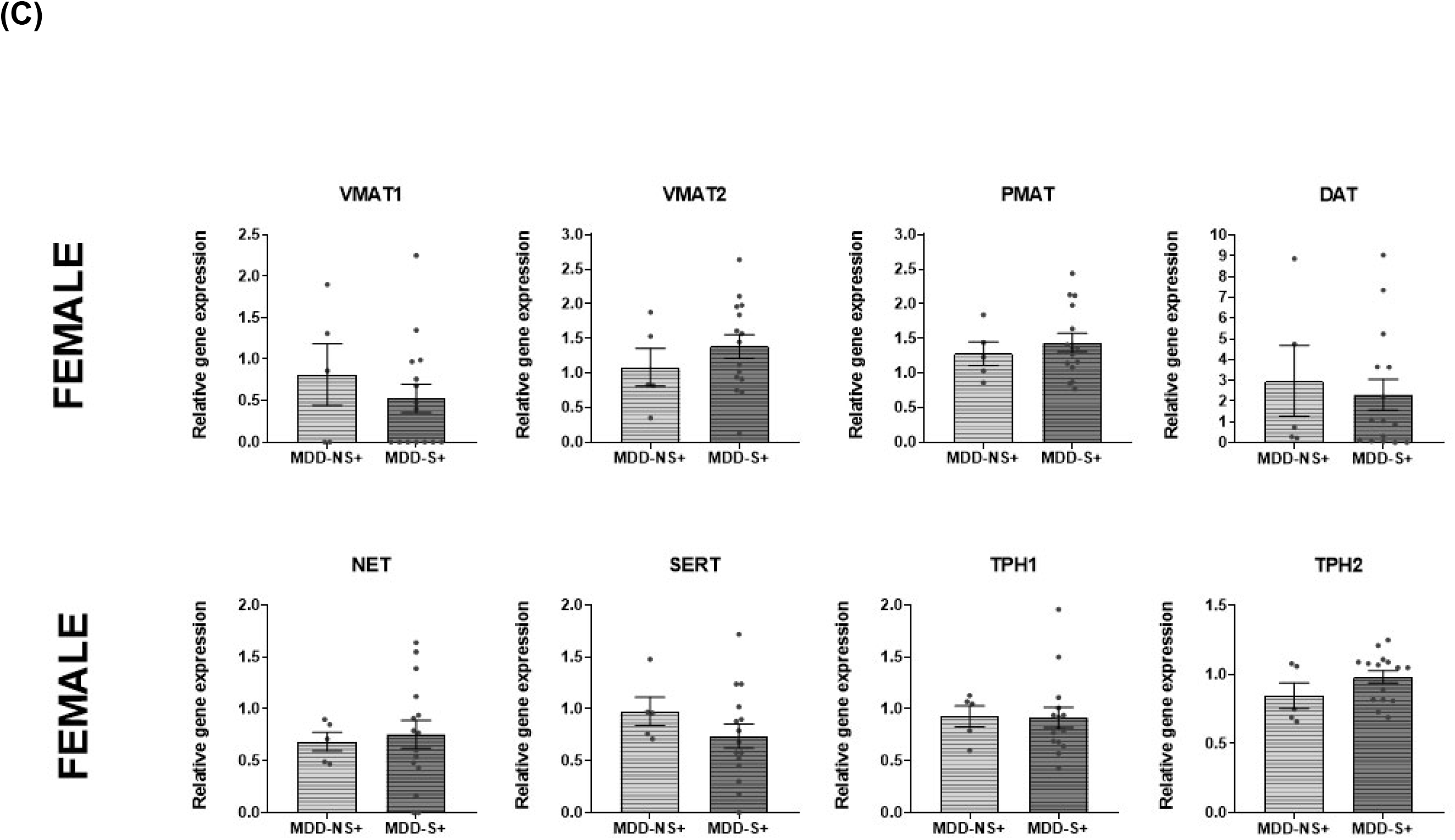
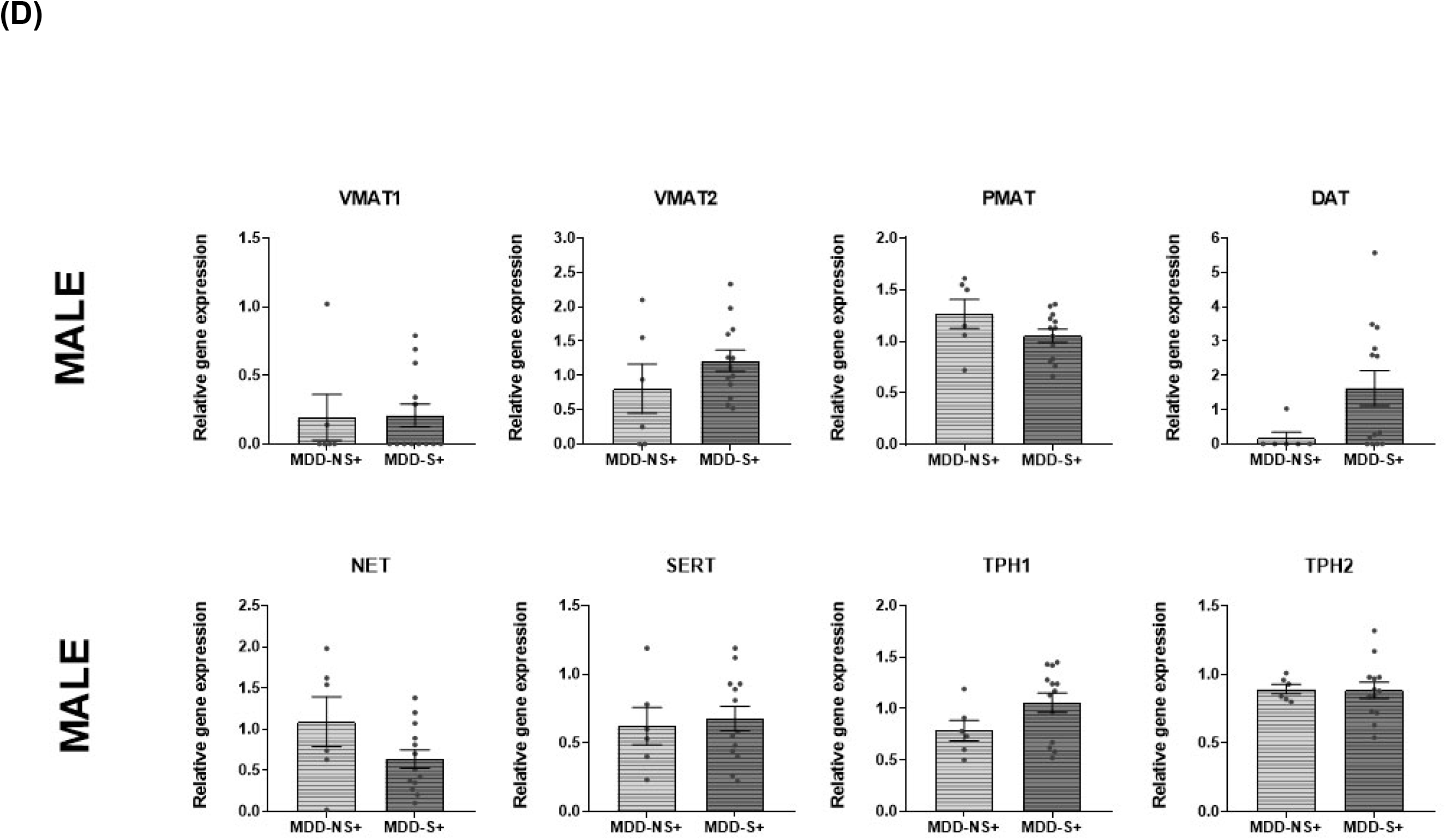
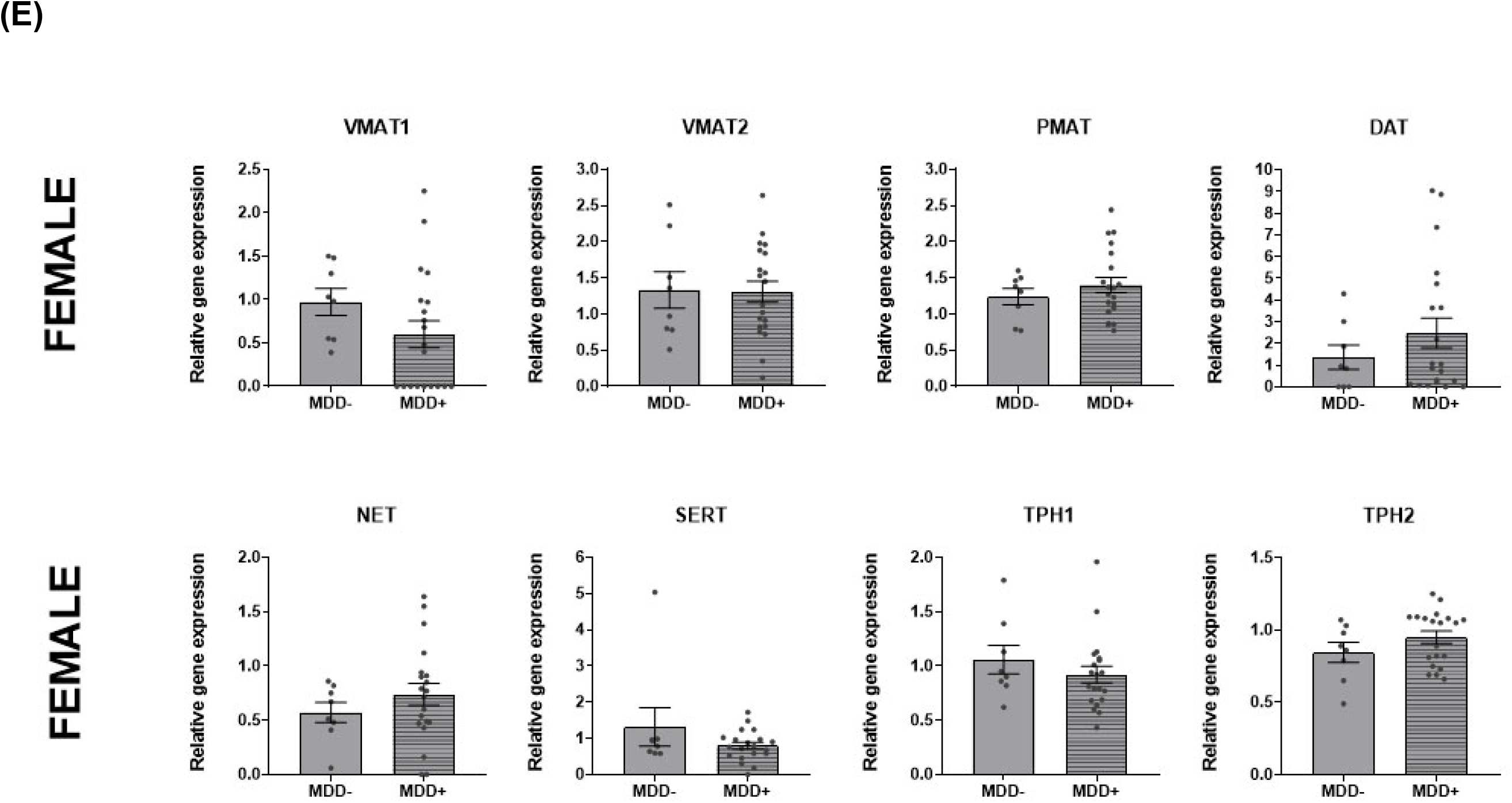
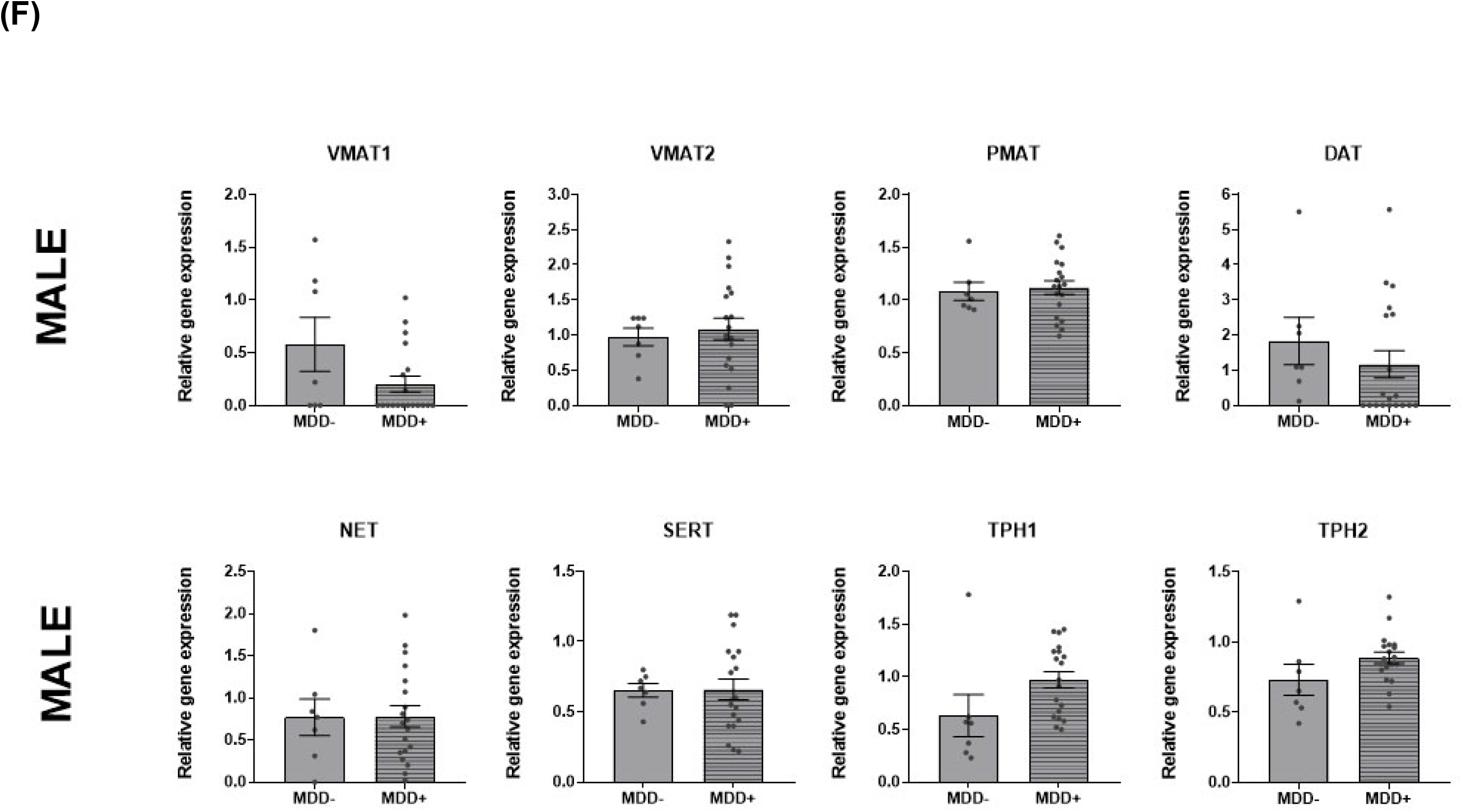
No association of monoaminergic gene expression in the DLPFC with suicide or antidepressant status. (A) and (B) No differences in monoaminergic gene expression between suicides and non-suicides with depression in males or females. (C) and (D) Suicide in subjects who were antidepressant positive at their time of death may be an indicator of antidepressant treatment response. Tests of depressed subjects who were antidepressant positive reveal no differences in the expression levels of monoaminergic genes between suicides and non-suicides, in males or females. (E) and (F). The monoaminergic proteins are targets of several conventional antidepressant drugs. Nevertheless, we detect no differences in the expression levels of monoaminergic genes between antidepressant positive and antidepressant negative depressed subjects, in either sex. *p<0.05, ** p<0.01.

**Supplementary Table 1.** Demographics of postmortem subjects.

| Diagnosis | Sex | N | Suicide<br>(Y / N) | Antidepressant<br>(+ / -) | Age<br>(mean ± sd) | PMI<br>(mean ± sd) | brain pH<br>(mean ± sd) | RIN<br>(mean ± sd) |
| --- | --- | --- | --- | --- | --- | --- | --- | --- |
| CTRL | M | 19 | 0 / 19 | 0 / 19 | 39.4 ± 12.7 | 30.6 ± 13.4 | 6.6 ± 0.3 | 8.3 ± 0.5 |
| | F | 13 | 0 / 13 | 0 / 13 | 38.9 ± 16.2 | 26.3 ± 12.0 | 6.4 ± 0.3 | 8.0 ± 1.1 |
| MDD | M | 39 | 15 / 24 | 19 / 7 | 41.9 ± 14.3 | 35.6 ± 27.3 | 6.4 ± 0.2 | 8.1 ± 0.8 |
| | F | 41 | 13 / 27 | 20 / 8 | 44.0 ± 14.8 | 35.8 ± 17.3 | 6.2 ± 0.3 | 8.1 ± 0.7 |

Abbreviations: Age at death in years (Age), antidepressant positive at time of death (+), antidepressant negative at time of death (-), Female (F), Male (M), number of subjects (N), Postmortem interval in hours (PMI), RNA integrity number (RIN), standard deviation (sd).

**Supplementary Table 2.** Gene expression assays tested in postmortem tissue.

| **Gene** | **Protein** | **Assay** |
| --- | --- | --- |
| SLC1A2 | Excitotoxic Amino Acid Transporter 2, EAAT2 | Hs001102423_m1 |
| SLC1A3 | Excitatory Amino Acid Transporter 1, EAAT1 | Hs00188193_m1 |
| SLC17A6 | Vesicular Glutamate Transporter 2, VGLUT2 | Hs00220439_m1 |
| SLC17A7 | Vesicular Glutamate Transporter 1, VGLUT1 | Hs00220404_m1 |
| SLC6A2 | Norepinephrine Transporter, NET | Hs00426573_m1 |
| SLC6A3 | Dopamine Transporter, DAT | Hs00997364_m1 |
| SLC6A4 | Serotonin Transporter, SERT | Hs00984349_m1 |
| SLC29A4 | Plasma Membrane Monoamine Transporter, PMAT | Hs00928283_m1 |
| SLC18A1 | Vesicular Amine Transporter 1, VMAT1 | Hs00915193_m1 |
| SLC18A2 | Vesicular Amine Transporter 2, VMAT2 | Hs00996835_m1 |
| TPH1 | Tryptophan hydroxylase 1 | Hs00188220_m1 |
| TPH2 | Tryptophan hydroxylase 2 | Hs00542783_m1 |
| ACTB | Beta-actin | Hs9999903_m1 |
| B2M | Beta-2-microglobulin | Hs9999907_m1 |
| GUSB | Beta-glucuronidase | Hs9999908_m1 |

**Supplementary Table 3.** Monoamine gene expression in the DLPFC does not differ between depressed suicides and non-suicides.

| Gene | Group | Sex | N | Mean | s.d. | Mean diff | statistic | p-value |
| --- | --- | --- | --- | --- | --- | --- | --- | --- |
| VMAT1 | MDD-NS | M | 15 | 0.61 | 0.73 | M 0.195 | U = 157<sup>†</sup> | *p* = 0.521 |
| | | F | 13 | 0.72 | 0.61 | F 0.076 | $F_{1,34} = 0.10$ | *p* = 0.758 |
|  | MDD-S | M | 24 | 0.37 | 0.45 |  |  |  |
|  |  | F | 27 | 0.61 | 0.66 |  |  |  |
| VMAT2 | MDD-NS | M | 15 | 1.10 | 0.76 | M -0.123 | $F_{1,33} = 0.27$ | *p* = 0.608 |
| | | F | 13 | 1.35 | 0.65 | F 0.008 | $F_{1,34} = 0.00$ | *p* = 0.975 |
|  | MDD-S | M | 24 | 1.21 | 0.62 |  |  |  |
|  |  | F | 27 | 1.47 | 0.65 |  |  |  |
| PMAT | MDD-NS | M | 15 | 1.13 | 0.28 | M 0.041 | $F_{1,33} = 0.17$ | *p* = 0.682 |
| | | F | 13 | 1.25 | 0.32 | F -0.128 | $F_{1,34} = 0.66$ | *p* = 0.422 |
|  | MDD-S | M | 24 | 1.09 | 0.31 |  |  |  |
|  |  | F | 27 | 1.39 | 0.32 |  |  |  |
| DAT | MDD-NS | M | 15 | 1.26 | 2.32 | M -0.284 | $F_{1,33} = 0.25$ | *p* = 0.621 |
| | | F | 13 | 1.76 | 2.56 | F -0.237 | $F_{1,34} = 0.07$ | *p* = 0.788 |
|  | MDD-S | M | 24 | 1.19 | 1.54 |  |  |  |
|  |  | F | 27 | 1.66 | 2.40 |  |  |  |
| NET | MDD-NS | M | 15 | 0.85 | 0.60 | M 0.209 | $F_{1,33} = 1.34$ | *p* = 0.255 |
|  |  | F | 13 | 0.73 | 0.21 | F 0.082 | U = 171<sup>†</sup> | *p* = 0.897 |
|  | MDD-S | M | 24 | 0.73 | 0.52 |  |  |  |
|  |  | F | 27 | 0.76 | 0.53 |  |  |  |
| SERT | MDD-NS | M | 15 | 0.68 | 0.28 | M -0.103 | $F_{1,33} = 0.90$ | *p* = 0.348 |
| | | F | 13 | 0.92 | 0.25 | F 0.107 | $F_{1,34} = 0.12$ | *p* = 0.727 |
|  | MDD-S | M | 24 | 0.76 | 0.35 |  |  |  |
|  |  | F | 27 | 0.94 | 0.96 |  |  |  |
| TPH1 | MDD-NS | M | 15 | 0.79 | 0.36 | M -0.207 | $F_{1,33} = 2.78$ | *p* = 0.105 |
| | | F | 13 | 0.95 | 0.30 | F -0.007 | $F_{1,34} = 0.00$ | *p* = 0.964 |
|  | MDD-S | M | 24 | 1.00 | 0.37 |  |  |  |
|  |  | F | 27 | 0.97 | 0.41 |  |  |  |
| TPH2 | MDD-NS | M | 15 | 0.81 | 0.15 | M -0.062 | U = 148<sup>†</sup> | *p* = 0.368 |
| | | F | 13 | 0.90 | 0.21 | F -0.043 | $F_{1,34} = 0.35$ | *p* = 0.558 |
|  | MDD-S | M | 24 | 0.90 | 0.30 |  |  |  |
|  |  | F | 27 | 0.96 | 0.20 |  |  |  |

Abbreviations: Antidepressant positive at time of death (+), Control (CTRL), standard deviation (sd), Major Depressive Disorder (MDD), number of subjects (N), probability (p). Female (F), Male (M), Major Depressive Disorder Non-Suicide (MDD-NS), Major Depressive Disorder Suicide (MDD-S), <sup>†</sup>Mann-Whitney U Test. Gene names listed in Supplementary Table 1.
<sup>†</sup>Mann-Whitney U Test, Gene names are listed in Supplementary Table 1.

**Supplementary Table 4.** Monoamine gene expression in the DLPFC does not differ between MDD-S and MDD-NS who were antidepressant positive at time of death by suicide.

| Gene | Group | Sex | N | Mean | s.d. |  | Mean diff | statistic | p-value |
| --- | --- | --- | --- | --- | --- | --- | --- | --- | --- |
| VMAT1 | MDD-NS+ | M | 6 | 0.19 | 0.41 | M | -0.278 | $F_{1,13} = 2.39$ | $p = 0.146$ |
| | | F | 5 | 0.81 | 0.83 | F | 0.108 | $F_{1,14} = 0.05$ | $p = 0.829$ |
|  | MDD-S+ | M | 13 | 0.21 | 0.30 |  |  |  |  |
|  |  | F | 15 | 0.52 | 0.66 |  |  |  |  |
| VMAT2 | MDD-NS+ | M | 6 | 0.81 | 0.88 | M | -0.167 | $F_{1,13} = 0.17$ | $p = 0.688$ |
| | | F | 5 | 1.08 | 0.61 | F | 0.251 | $F_{1,14} = 0.30$ | $p = 0.595$ |
|  | MDD-S+ | M | 13 | 1.21 | 0.55 |  |  |  |  |
|  |  | F | 15 | 1.38 | 0.66 |  |  |  |  |
| PMAT | MDD-NS+ | M | 6 | 1.26 | 0.35 | M | 0.263 | $F_{1,13} = 2.55$ | $p = 0.134$ |
| | | F | 5 | 1.28 | 0.38 | F | 0.218 | $F_{1,14} = 0.54$ | $p = 0.474$ |
|  | MDD-S+ | M | 13 | 1.05 | 0.23 |  |  |  |  |
|  |  | F | 15 | 1.44 | 0.52 |  |  |  |  |
| DAT | MDD-NS+ | M | 6 | 0.17 | 0.42 | M | -1.455 | $U = 17^{\dagger}$ | $p = 0.058$ |
| | | F | 5 | 2.97 | 3.80 | F | 0.502 | $F_{1,14} = 0.04$ | $p = 0.841$ |
|  | MDD-S+ | M | 13 | 1.63 | 1.86 |  |  |  |  |
|  |  | F | 15 | 2.31 | 2.90 |  |  |  |  |
| NET | MDD-NS+ | M | 6 | 1.09 | 0.74 | M | 0.654 | $F_{1,13} = 4.62$ | $p = 0.051$ |
| | | F | 5 | 0.68 | 0.20 | F | 0.178 | $F_{1,14} = 0.27$ | $p = 0.608$ |
|  | MDD-S+ | M | 13 | 0.64 | 0.41 |  |  |  |  |
|  |  | F | 15 | 0.75 | 0.52 |  |  |  |  |
| SERT | MDD-NS+ | M | 6 | 0.62 | 0.33 | M | 0.135 | $F_{1,13} = 0.52$ | $p = 0.483$ |
| | | F | 5 | 0.97 | 0.31 | F | 0.322 | $F_{1,14} = 1.32$ | $p = 0.269$ |
|  | MDD-S+ | M | 13 | 0.68 | 0.32 |  |  |  |  |
|  |  | F | 15 | 0.74 | 0.45 |  |  |  |  |
| TPH1 | MDD-NS+ | M | 6 | 0.78 | 0.24 | M | -0.227 | $F_{1,13} = 1.11$ | $p = 0.312$ |
| | | F | 5 | 0.93 | 0.23 | F | -0.079 | $F_{1,14} = 0.08$ | $p = 0.781$ |
|  | MDD-S+ | M | 13 | 1.05 | 0.34 |  |  |  |  |
|  |  | F | 15 | 0.92 | 0.38 |  |  |  |  |
| TPH2 | MDD-NS+ | M | 6 | 0.89 | 0.84 | M | 0.031 | $F_{1,13} = 0.06$ | $p = 0.805$ |
| | | F | 5 | 0.85 | 0.20 | F | -0.144 | $F_{1,14} = 2.16$ | $p = 0.164$ |
|  | MDD-S+ | M | 13 | 0.88 | 0.21 |  |  |  |  |
|  |  | F | 15 | 0.98 | 0.18 |  |  |  |  |

Abbreviations: Antidepressant positive at time of death (+), Control (CTRL), standard deviation (sd), Major Depressive Disorder (MDD), number of subjects (N), probability (p). Female (F), Male (M), Major Depressive Disorder Non-Suicide (MDD-NS), Major Depressive Disorder Suicide (MDD-S), $^{\dagger}$ Mann-Whitney U Test. Gene names listed in Supplementary Table 1.

**Supplementary Table 5.** Monoamine gene expression in the DLPFC does not differ between depressed subjects who were antidepressant positive and antidepressant negative.

| Gene | Group | Sex | N | Mean | s.d. |  | Mean diff | statistic | p-value |
| --- | --- | --- | --- | --- | --- | --- | --- | --- | --- |
| VMAT1 | MDD- | M | 7 | 0.58 | 0.67 | M | 0.355 | $F_{1,20} = 2.82$ | $p = 0.108$ |
| | | F | 8 | 0.97 | 0.44 | F | 0.388 | $F_{1,22} = 1.68$ | $p = 0.208$ |
|  | MDD+ | M | 19 | 0.20 | 0.33 |  |  |  |  |
|  |  | F | 20 | 0.60 | 0.69 |  |  |  |  |
| VMAT2 | MDD- | M | 7 | 0.97 | 0.33 | M | -0.158 | $F_{1,20} = 0.25$ | $p = 0.619$ |
| | | F | 8 | 1.33 | 0.72 | F | 0.077 | $F_{1,22} = 0.07$ | $p = 0.788$ |
|  | MDD+ | M | 19 | 1.08 | 0.67 |  |  |  |  |
|  |  | F | 20 | 1.31 | 0.65 |  |  |  |  |
| PMAT | MDD- | M | 7 | 1.08 | 0.23 | M | -0.045 | $F_{1,20} = 0.11$ | $p = 0.745$ |
| | | F | 8 | 1.24 | 0.32 | F | -0.172 | $F_{1,22} = 0.84$ | $p = 0.370$ |
|  | MDD+ | M | 19 | 1.12 | 0.28 |  |  |  |  |
|  |  | F | 20 | 1.40 | 0.48 |  |  |  |  |
| DAT | MDD- | M | 7 | 1.83 | 1.78 | M | -0.195 | $F_{1,20} = 0.06$ | $p = 0.801$ |
| | | F | 8 | 1.37 | 1.58 | F | -0.871 | $U = 62^{\dagger}$ | $p = 0.359$ |
|  | MDD+ | M | 19 | 1.17 | 1.68 |  |  |  |  |
|  |  | F | 20 | 2.47 | 3.05 |  |  |  |  |
| NET | MDD- | M | 7 | 0.77 | 0.57 | M | 0.183 | $F_{1,20} = 0.45$ | $p = 0.511$ |
| | | F | 8 | 0.57 | 0.26 | F | -0.054 | $U = 62^{\dagger}$ | $p = 0.360$ |
|  | MDD+ | M | 19 | 0.78 | 0.56 |  |  |  |  |
|  |  | F | 20 | 0.73 | 0.45 |  |  |  |  |
| SERT | MDD- | M | 7 | 0.65 | 0.13 | M | -0.071 | $U = 63^{\dagger}$ | $p = 0.840$ |
| | | F | 8 | 1.31 | 1.51 | F | 0.472 | $U = 68^{\dagger}$ | $p = 0.542$ |
|  | MDD+ | M | 19 | 0.66 | 0.32 |  |  |  |  |
|  |  | F | 20 | 0.80 | 0.42 |  |  |  |  |
| TPH1 | MDD- | M | 7 | 0.63 | 0.53 | M | -0.295 | $F_{1,20} = 2.31$ | $p = 0.144$ |
| | | F | 8 | 1.06 | 0.37 | F | 0.066 | $F_{1,22} = 0.14$ | $p = 0.707$ |
|  | MDD+ | M | 19 | 0.97 | 0.33 |  |  |  |  |
|  |  | F | 20 | 0.92 | 0.34 |  |  |  |  |
| TPH2 | MDD- | M | 7 | 0.73 | 0.29 | M | -0.095 | $F_{1,20} = 0.96$ | $p = 0.339$ |
| | | F | 8 | 0.85 | 0.20 | F | -0.022 | $F_{1,22} = 0.08$ | $p = 0.779$ |
|  | MDD+ | M | 19 | 0.89 | 0.18 |  |  |  |  |
|  |  | F | 20 | 0.95 | 0.19 |  |  |  |  |

Abbreviations: Antidepressant positive at time of death (+), antidepressant negative at time of death (-), Control (CTRL), standard deviation (sd), Major Depressive Disorder (MDD), number of subjects (N), probability (p). Female (F), Male (M), Non-Suicide (NS), Suicide (S), $^{\dagger}$ Mann-Whitney U Test. Gene names listed in Supplementary Table 1.

## References

1. Cavanagh JT, Carson AJ, Sharpe M, Lawrie SM. Psychological autopsy studies of suicide: a systematic review. Psychol Med. 2003;33(3):395–405.

2. Turecki G, Brent DA. Suicide and suicidal behaviour. Lancet. 2016;387(10024):1227–39. doi:10.1016/S0140-6736(15)00234-2

3. Charney DS. Monoamine dysfunction and the pathophysiology and treatment of depression. J Clin Psychiatry. 1998;59 Suppl 14:11–4.

4. Duric V, Duman RS. Depression and treatment response: dynamic interplay of signaling pathways and altered neural processes. Cell Mol Life Sci. 2013;70(1):39–53. doi:10.1007/s00018-012-1020-7

5. Shelton RC. Serotonin and Norepinephrine Reuptake Inhibitors. Handb Exp Pharmacol. 2019. doi:10.1007/164_2018_164

6. Belmaker RH, Agam G. Major depressive disorder. N Engl J Med. 2008;358(1):55–68. doi:10.1056/NEJMra073096

7. Trivedi MH, Rush AJ, Wisniewski SR, Nierenberg AA, Warden D, Ritz L, et al. Evaluation of outcomes with citalopram for depression using measurement-based care in STAR*D: implications for clinical practice. Am J Psychiatry. 2006;163(1):28–40. doi:10.1176/appi.ajp.163.1.28

8. Block SG, Nemeroff CB. Emerging antidepressants to treat major depressive disorder. Asian J Psychiatr. 2014;12:7–16. doi:10.1016/j.ajp.2014.09.001

9. Cai S, Huang S, Hao W. New hypothesis and treatment targets of depression: an integrated view of key findings. Neurosci Bull. 2015;31(1):61–74. doi:10.1007/s12264-014-1486-4

10. Peng GJ, Tian JS, Gao XX, Zhou YZ, Qin XM. Research on the Pathological Mechanism and Drug Treatment Mechanism of Depression. Curr Neuropharmacol. 2015;13(4):514–23.

11. Berman RM, Cappiello A, Anand A, Oren DA, Heninger GR, Charney DS, et al. Antidepressant effects of ketamine in depressed patients. Biol Psychiatry. 2000;47(4):351–4.

12. Sanacora G, Treccani G, Popoli M. Towards a glutamate hypothesis of depression: an emerging frontier of neuropsychopharmacology for mood disorders. Neuropharmacology. 2012;62(1):63–77. doi:10.1016/j.neuropharm.2011.07.036

13. Zarate C, Jr., Machado-Vieira R, Henter I, Ibrahim L, Diazgranados N, Salvadore G. Glutamatergic modulators: the future of treating mood disorders? Harv Rev Psychiatry. 2010;18(5):293–303. doi:10.3109/10673229.2010.511059

14. Abdallah CG, Sanacora G, Duman RS, Krystal JH. The neurobiology of depression, ketamine and rapid-acting antidepressants: Is it glutamate inhibition or activation? Pharmacol Ther. 2018;190:148–58. doi:10.1016/j.pharmthera.2018.05.010

15. Duman RS. Ketamine and rapid-acting antidepressants: a new era in the battle against depression and suicide. F1000Res. 2018;7. doi:10.12688/f1000research.14344.1

16. Kadriu B, Musazzi L, Henter ID, Graves M, Popoli M, Zarate CA, Jr. Glutamatergic Neurotransmission: Pathway to Developing Novel Rapid-Acting Antidepressant Treatments. Int J Neuropsychopharmacol. 2019;22(2):119–35. doi:10.1093/ijnp/pyy094

17. Kessler RC, Chiu WT, Demler O, Merikangas KR, Walters EE. Prevalence, severity, and comorbidity of 12-month DSM-IV disorders in the National Comorbidity Survey Replication. Arch Gen Psychiatry. 2005;62(6):617–27. doi:10.1001/archpsyc.62.6.617

18. Kuehner C. Gender differences in unipolar depression: an update of epidemiological findings and possible explanations. Acta Psychiatr Scand. 2003;108(3):163–74.

19. Kessler RC. Epidemiology of women and depression. J Affect Disord. 2003;74(1):5–13.

20. Kornstein SG, Schatzberg AF, Thase ME, Yonkers KA, McCullough JP, Keitner GI, et al. Gender differences in chronic major and double depression. J Affect Disord. 2000;60(1):1–11.

21. Labonte B, Engmann O, Purushothaman I, Menard C, Wang J, Tan C, et al. Sex-specific transcriptional signatures in human depression. Nat Med. 2017;23(9):1102–11. doi:10.1038/nm.4386

22. Seney ML, Huo Z, Cahill K, French L, Puralewski R, Zhang J, et al. Opposite Molecular Signatures of Depression in Men and Women. Biol Psychiatry. 2018;84(1):18–27. doi:10.1016/j.biopsych.2018.01.017

23. Goldstein JM, Hale T, Foster SL, Tobet SA, Handa RJ. Sex differences in major depression and comorbidity of cardiometabolic disorders: impact of prenatal stress and immune exposures. Neuropsychopharmacology. 2019;44(1):59–70. doi:10.1038/s41386-018-0146-1

24. Levy R, Goldman-Rakic PS. Segregation of working memory functions within the dorsolateral prefrontal cortex. Exp Brain Res. 2000;133(1):23–32.

25. Dolcos F, Diaz-Granados P, Wang L, McCarthy G. Opposing influences of emotional and non-emotional distracters upon sustained prefrontal cortex activity during a delayed-response working memory task. Neuropsychologia. 2008;46(1):326–35. doi:S0028-3932(07)00249-7 [pii] 10.1016/j.neuropsychologia.2007.07.010

26. Dolcos F, McCarthy G. Brain systems mediating cognitive interference by emotional distraction. The Journal of neuroscience : the official journal of the Society for Neuroscience. 2006;26(7):2072–9. doi:26/7/2072 [pii] 10.1523/JNEUROSCI.5042-05.2006

27. Perlstein WM, Elbert T, Stenger VA. Dissociation in human prefrontal cortex of affective influences on working memory-related activity. Proceedings of the National Academy of Sciences of the United States of America. 2002;99(3):1736–41. doi:10.1073/pnas.241650598 241650598 [pii]

28. Byrum CE, Ahearn EP, Krishnan KR. A neuroanatomic model for depression. Prog Neuropsychopharmacol Biol Psychiatry. 1999;23(2):175–93.

29. Fitzgerald PB, Oxley TJ, Laird AR, Kulkarni J, Egan GF, Daskalakis ZJ. An analysis of functional neuroimaging studies of dorsolateral prefrontal cortical activity in depression. Psychiatry Res. 2006;148(1):33–45. doi:S0925-4927(06)00068-0 [pii] 10.1016/j.pscychresns.2006.04.006

30. Kang HJ, Voleti B, Hajszan T, Rajkowska G, Stockmeier CA, Licznerski P, et al. Decreased expression of synapse-related genes and loss of synapses in major depressive disorder. Nat Med. 2012;18(9):1413–7. doi:nm.2886 [pii] 10.1038/nm.2886

31. Holtzheimer PE, Kelley ME, Gross RE, Filkowski MM, Garlow SJ, Barrocas A, et al. Subcallosal cingulate deep brain stimulation for treatment-resistant unipolar and bipolar depression. Arch Gen Psychiatry. 2012;69(2):150–8. doi:10.1001/archgenpsychiatry.2011.1456 archgenpsychiatry.2011.1456 [pii]

32. Kennedy SH, Giacobbe P, Rizvi SJ, Placenza FM, Nishikawa Y, Mayberg HS, et al. Deep brain stimulation for treatment-resistant depression: follow-up after 3 to 6 years. The American journal of psychiatry. 2011;168(5):502–10. doi:10.1176/appi.ajp.2010.10081187 appi.ajp.2010.10081187 [pii]

33. Lozano AM, Mayberg HS, Giacobbe P, Hamani C, Craddock RC, Kennedy SH. Subcallosal cingulate gyrus deep brain stimulation for treatment-resistant depression. Biological psychiatry. 2008;64(6):461–7. doi:10.1016/j.biopsych.2008.05.034 S0006-3223(08)00703-8 [pii]

34. Mayberg HS, Lozano AM, Voon V, McNeely HE, Seminowicz D, Hamani C, et al. Deep brain stimulation for treatment-resistant depression. Neuron. 2005;45(5):651–60. doi:S0896-6273(05)00156-X [pii] 10.1016/j.neuron.2005.02.014

35. Fox MD, Buckner RL, White MP, Greicius MD, Pascual-Leone A. Efficacy of transcranial magnetic stimulation targets for depression is related to intrinsic functional connectivity with the subgenual cingulate. Biol Psychiatry. 2012;72(7):595–603. doi:10.1016/j.biopsych.2012.04.028 S0006-3223(12)00411-8 [pii]

36. Fox MD, Liu H, Pascual-Leone A. Identification of reproducible individualized targets for treatment of depression with TMS based on intrinsic connectivity. Neuroimage. 2013;66:151–60. doi:10.1016/j.neuroimage.2012.10.082 S1053-8119(12)01086-5 [pii]

37. Duman RS. Pathophysiology of depression and innovative treatments: remodeling glutamatergic synaptic connections. Dialogues Clin Neurosci. 2014;16(1):11–27.

38. Choudary PV, Molnar M, Evans SJ, Tomita H, Li JZ, Vawter MP, et al. Altered cortical glutamatergic and GABAergic signal transmission with glial involvement in depression. Proc Natl Acad Sci U S A. 2005;102(43):15653–8. doi:10.1073/pnas.0507901102

39. Gray AL, Hyde TM, Deep-Soboslay A, Kleinman JE, Sodhi MS. Sex differences in glutamate receptor gene expression in major depression and suicide. Mol Psychiatry. 2015;20(9):1057–68. doi:10.1038/mp.2015.91

40. Zhao J, Verwer RWH, Gao SF, Qi XR, Lucassen PJ, Kessels HW, et al. Prefrontal alterations in GABAergic and glutamatergic gene expression in relation to depression and suicide. J Psychiatr Res. 2018;102:261–74. doi:10.1016/j.jpsychires.2018.04.020

41. Arango V, Underwood MD, Mann JJ. Serotonin brain circuits involved in major depression and suicide. Prog Brain Res. 2002;136:443–53.

42. Charney DS, Krystal JH, Delgado PL, Heninger GR. Serotonin-specific drugs for anxiety and depressive disorders. Annu Rev Med. 1990;41:437–46. doi:10.1146/annurev.me.41.020190.002253

43. Sodhi MS, Sanders-Bush E. Serotonin and brain development. Int Rev Neurobiol. 2004;59:111–74. doi:10.1016/S0074-7742(04)59006-2

44. Mann JJ, Huang YY, Underwood MD, Kassir SA, Oppenheim S, Kelly TM, et al. A serotonin transporter gene promoter polymorphism (5-HTTLPR) and prefrontal cortical binding in major depression and suicide. Arch Gen Psychiatry. 2000;57(8):729–38.

45. Klimek V, Stockmeier C, Overholser J, Meltzer HY, Kalka S, Dilley G, et al. Reduced levels of norepinephrine transporters in the locus coeruleus in major depression. J Neurosci. 1997;17(21):8451–8.

46. Nutt DJ. The role of dopamine and norepinephrine in depression and antidepressant treatment. J Clin Psychiatry. 2006;67 Suppl 6:3–8.

47. Santen G, Gomeni R, Danhof M, Della Pasqua O. Sensitivity of the individual items of the Hamilton depression rating scale to response and its consequences for the assessment of efficacy. J Psychiatr Res. 2008;42(12):1000–9. doi:10.1016/j.jpsychires.2007.11.004

48. Rajkowska G, Goldman-Rakic PS. Cytoarchitectonic definition of prefrontal areas in the normal human cortex: I. Remapping of areas 9 and 46 using quantitative criteria. Cereb Cortex. 1995;5(4):307–22.

49. Mengual L, Burset M, Marin-Aguilera M, Ribal MJ, Alcaraz A. Multiplex preamplification of specific cDNA targets prior to gene expression analysis by TaqMan Arrays. BMC Res Notes. 2008;1:21. doi:10.1186/1756-0500-1-21

50. Sodhi MS, Simmons M, McCullumsmith R, Haroutunian V, Meador-Woodruff JH. Glutamatergic gene expression is specifically reduced in thalamocortical projecting relay neurons in schizophrenia. Biol Psychiatry. 2011;70(7):646–54. doi:S0006-3223(11)00197-1 [pii]10.1016/j.biopsych.2011.02.022

51. Benjamini Y, Hochberg Y. Controlling the False Discovery Rate - a Practical and Powerful Approach to Multiple Testing. J R Stat Soc B. 1995;57(1):289–300.

52. Hashimoto K, Sawa A, Iyo M. Increased levels of glutamate in brains from patients with mood disorders. Biol Psychiatry. 2007;62(11):1310–6. doi:10.1016/j.biopsych.2007.03.017

53. Yuen EY, Wei J, Yan Z. Estrogen in prefrontal cortex blocks stress-induced cognitive impairments in female rats. J Steroid Biochem Mol Biol. 2016;160:221–6. doi:10.1016/j.jsbmb.2015.08.028

54. Bourdeau V, Deschenes J, Metivier R, Nagai Y, Nguyen D, Bretschneider N, et al. Genome-wide identification of high-affinity estrogen response elements in human and mouse. Mol Endocrinol. 2004;18(6):1411–27. doi:10.1210/me.2003-0441

55. Tian G, Lai L, Guo H, Lin Y, Butchbach ME, Chang Y, et al. Translational control of glial glutamate transporter EAAT2 expression. J Biol Chem. 2007;282(3):1727–37. doi:10.1074/jbc.M609822200

56. Zschocke J, Bayatti N, Clement AM, Witan H, Figiel M, Engele J, et al. Differential promotion of glutamate transporter expression and function by glucocorticoids in astrocytes from various brain regions. J Biol Chem. 2005;280(41):34924–32. doi:10.1074/jbc.M502581200

57. Oh DH, Oh D, Son H, Webster MJ, Weickert CS, Kim SH. An association between the reduced levels of SLC1A2 and GAD1 in the dorsolateral prefrontal cortex in major depressive disorder: possible involvement of an attenuated RAF/MEK/ERK signaling pathway. J Neural Transm (Vienna). 2014;121(7):783–92. doi:10.1007/s00702-014-1189-z

58. Zhao J, Verwer RW, van Wamelen DJ, Qi XR, Gao SF, Lucassen PJ, et al. Prefrontal changes in the glutamate-glutamine cycle and neuronal/glial glutamate transporters in depression with and without suicide. J Psychiatr Res. 2016;82:8–15. doi:10.1016/j.jpsychires.2016.06.017

59. Miguel-Hidalgo JJ, Waltzer R, Whittom AA, Austin MC, Rajkowska G, Stockmeier CA. Glial and glutamatergic markers in depression, alcoholism, and their comorbidity. J Affect Disord. 2010;127(1-3):230–40. doi:10.1016/j.jad.2010.06.003

60. Gilabert-Juan J, Varea E, Guirado R, Blasco-Ibanez JM, Crespo C, Nacher J. Alterations in the expression of PSA-NCAM and synaptic proteins in the dorsolateral prefrontal cortex of psychiatric disorder patients. Neurosci Lett. 2012;530(1):97–102. doi:10.1016/j.neulet.2012.09.032

61. Shao L, Vawter MP. Shared gene expression alterations in schizophrenia and bipolar disorder. Biol Psychiatry. 2008;64(2):89–97. doi:10.1016/j.biopsych.2007.11.010

62. Medina A, Burke S, Thompson RC, Bunney W, Jr., Myers RM, Schatzberg A, et al. Glutamate transporters: a key piece in the glutamate puzzle of major depressive disorder. J Psychiatr Res. 2013;47(9):1150–6. doi:10.1016/j.jpsychires.2013.04.007

63. Chandley MJ, Szebeni K, Szebeni A, Crawford J, Stockmeier CA, Turecki G, et al. Gene expression deficits in pontine locus coeruleus astrocytes in men with major depressive disorder. J Psychiatry Neurosci. 2013;38(4):276–84. doi:10.1503/jpn.120110

64. Roberts RC, Roche JK, McCullumsmith RE. Localization of excitatory amino acid transporters EAAT1 and EAAT2 in human postmortem cortex: a light and electron microscopic study. Neuroscience. 2014;277:522–40. doi:10.1016/j.neuroscience.2014.07.019

65. O’Donovan SM, Sullivan CR, McCullumsmith RE. The role of glutamate transporters in the pathophysiology of neuropsychiatric disorders. NPJ Schizophr. 2017;3(1):32. doi:10.1038/s41537-017-0037-1

66. Parkin GM, Udawela M, Gibbons A, Dean B. Glutamate transporters, EAAT1 and EAAT2, are potentially important in the pathophysiology and treatment of schizophrenia and affective disorders. World J Psychiatry. 2018;8(2):51–63. doi:10.5498/wjp.v8.i2.51

67. van Heeringen C, Bijttebier S, Godfrin K. Suicidal brains: a review of functional and structural brain studies in association with suicidal behaviour. Neurosci Biobehav Rev. 2011;35(3):688–98. doi:10.1016/j.neubiorev.2010.08.007

68. Grimm S, Luborzewski A, Schubert F, Merkl A, Kronenberg G, Colla M, et al. Region-specific glutamate changes in patients with unipolar depression. J Psychiatr Res. 2012;46(8):1059–65. doi:10.1016/j.jpsychires.2012.04.018

69. Dossat AM, Wright KN, Strong CE, Kabbaj M. Behavioral and biochemical sensitivity to low doses of ketamine: Influence of estrous cycle in C57BL/6 mice. Neuropharmacology. 2018;130:30–41. doi:10.1016/j.neuropharm.2017.11.022

70. Cowen PJ, Browning M. What has serotonin to do with depression? World Psychiatry. 2015;14(2):158–60. doi:10.1002/wps.20229

71. Kristensen AS, Andersen J, Jorgensen TN, Sorensen L, Eriksen J, Loland CJ, et al. SLC6 neurotransmitter transporters: structure, function, and regulation. Pharmacol Rev. 2011;63(3):585–640. doi:10.1124/pr.108.000869

72. Wang J. The plasma membrane monoamine transporter (PMAT): Structure, function, and role in organic cation disposition. Clin Pharmacol Ther. 2016;100(5):489–99. doi:10.1002/cpt.442

73. Steinkellner T, Zell V, Farino ZJ, Sonders MS, Villeneuve M, Freyberg RJ, et al. Role for VGLUT2 in selective vulnerability of midbrain dopamine neurons. J Clin Invest. 2018;128(2):774–88. doi:10.1172/JCI95795

74. Perroud N, Neidhart E, Petit B, Vessaz M, Laforge T, Relecom C, et al. Simultaneous analysis of serotonin transporter, tryptophan hydroxylase 1 and 2 gene expression in the ventral prefrontal cortex of suicide victims. Am J Med Genet B Neuropsychiatr Genet. 2010;153B(4):909–18. doi:10.1002/ajmg.b.31059

75. De Luca V, Likhodi O, Van Tol HH, Kennedy JL, Wong AH. Gene expression of tryptophan hydroxylase 2 in post-mortem brain of suicide subjects. Int J Neuropsychopharmacol. 2006;9(1):21–5. doi:10.1017/S1461145705005572

76. Kambeitz JP, Howes OD. The serotonin transporter in depression: Meta-analysis of in vivo and post mortem findings and implications for understanding and treating depression. J Affect Disord. 2015;186:358–66. doi:10.1016/j.jad.2015.07.034

77. Deep-Soboslay A, Benes FM, Haroutunian V, Ellis JK, Kleinman JE, Hyde TM. Psychiatric brain banking: three perspectives on current trends and future directions. Biol Psychiatry. 2011;69(2):104–12. doi:10.1016/j.biopsych.2010.05.025 S0006-3223(10)00498-1 [pii]

78. Amrhein V, Greenland S, McShane B. Scientists rise up against statistical significance. Nature. 2019;567(7748):305–7. doi:10.1038/d41586-019-00857-9

79. Grassi S, Frondaroli A, Scarduzio M, Dutia MB, Dieni C, Pettorossi VE. Effects of 17beta-estradiol on glutamate synaptic transmission and neuronal excitability in the rat medial vestibular nuclei. Neuroscience. 2010;165(4):1100–14. doi:10.1016/j.neuroscience.2009.11.039

80. Nebieridze N, Zhang XL, Chachua T, Velisek L, Stanton PK, Veliskova J. beta-Estradiol unmasks metabotropic receptor-mediated metaplasticity of NMDA receptor transmission in the female rat dentate gyrus. Psychoneuroendocrinology. 2012;37(11):1845–54. doi:10.1016/j.psyneuen.2012.03.023

81. Rothstein JD, Dykes-Hoberg M, Pardo CA, Bristol LA, Jin L, Kuncl RW, et al. Knockout of glutamate transporters reveals a major role for astroglial transport in excitotoxicity and clearance of glutamate. Neuron. 1996;16(3):675–86.

82. Tzingounis AV, Wadiche JI. Glutamate transporters: confining runaway excitation by shaping synaptic transmission. Nat Rev Neurosci. 2007;8(12):935–47. doi:10.1038/nrn2274

83. Valentine GW, Sanacora G. Targeting glial physiology and glutamate cycling in the treatment of depression. Biochem Pharmacol. 2009;78(5):431–9. doi:10.1016/j.bcp.2009.04.008

84. Cotter DR, Pariante CM, Everall IP. Glial cell abnormalities in major psychiatric disorders: the evidence and implications. Brain Res Bull. 2001;55(5):585–95.

85. Ongur D, Drevets WC, Price JL. Glial reduction in the subgenual prefrontal cortex in mood disorders. Proc Natl Acad Sci U S A. 1998;95(22):13290–5.

86. Rajkowska G, Halaris A, Selemon LD. Reductions in neuronal and glial density characterize the dorsolateral prefrontal cortex in bipolar disorder. Biol Psychiatry. 2001;49(9):741–52.

87. Rajkowska G, Miguel-Hidalgo JJ, Wei J, Dilley G, Pittman SD, Meltzer HY, et al. Morphometric evidence for neuronal and glial prefrontal cell pathology in major depression. Biol Psychiatry. 1999;45(9):1085–98.

88. Akiyama T, Koeda M, Okubo Y, Kimura M. Hypofunction of left dorsolateral prefrontal cortex in depression during verbal fluency task: A multi-channel near-infrared spectroscopy study. J Affect Disord. 2018;231:83–90. doi:10.1016/j.jad.2018.01.010

89. Korgaonkar MS, Grieve SM, Etkin A, Koslow SH, Williams LM. Using standardized fMRI protocols to identify patterns of prefrontal circuit dysregulation that are common and specific to cognitive and emotional tasks in major depressive disorder: first wave results from the iSPOT-D study. Neuropsychopharmacology. 2013;38(5):863–71. doi:10.1038/npp.2012.252

90. Liu X, Sun G, Zhang X, Xu B, Shen C, Shi L, et al. Relationship between the prefrontal function and the severity of the emotional symptoms during a verbal fluency task in patients with major depressive disorder: a multi-channel NIRS study. Prog Neuropsychopharmacol Biol Psychiatry. 2014;54:114–21. doi:10.1016/j.pnpbp.2014.05.005

91. Opel N, Redlich R, Grotegerd D, Dohm K, Zaremba D, Meinert S, et al. Prefrontal brain responsiveness to negative stimuli distinguishes familial risk for major depression from acute disorder. J Psychiatry Neurosci. 2017;42(5):343–52.

92. Schiller CE, Minkel J, Smoski MJ, Dichter GS. Remitted major depression is characterized by reduced prefrontal cortex reactivity to reward loss. J Affect Disord. 2013;151(2):756–62. doi:10.1016/j.jad.2013.06.016

93. Takamura M, Okamoto Y, Okada G, Toki S, Yamamoto T, Yamamoto O, et al. Disrupted Brain Activation and Deactivation Pattern during Semantic Verbal Fluency Task in Patients with Major Depression. Neuropsychobiology. 2016;74(2):69–77. doi:10.1159/000453399

94. Zhong M, Wang X, Xiao J, Yi J, Zhu X, Liao J, et al. Amygdala hyperactivation and prefrontal hypoactivation in subjects with cognitive vulnerability to depression. Biol Psychol. 2011;88(2-3):233–42. doi:10.1016/j.biopsycho.2011.08.007

